# Context-Dependent Protein Structure Prediction Analysis and Stoichiometry Inference with MultimerMapper

**DOI:** 10.1101/2025.10.24.684463

**Authors:** Elvio Rodríguez Araya, Esteban Serra

## Abstract

Advances in artificial intelligence (AI) have transformed the field of protein structure, notably with the accuracy level reached by AlphaFold in the prediction of monomeric and multimeric protein structures. However, while cloud implementations have broadened access to these methods, there is still a lack of non-AI tools to systematically interpret and analyze the resulting data, especially under varying modeling contexts. Here, we present MultimerMapper, a pipeline and software suite designed to extract, integrate, analyze, and visualize the behavior of multimeric systems in ensembles of overlapping protein structure predictions. It utilizes statistics and graph theory, combined with novel approaches, to interpret how protein-protein interfaces and protein conformation behave across different modeling contexts. Starting from a list of input sequences, MultimerMapper guides users through the generation and interpretation of structure prediction ensembles. It produces interactive 2D and 3D visualizations, highlights higher-order subassemblies, infers probable stoichiometries, and identifies alternative interaction modes and conformational changes associated with the presence or absence of specific proteins. MultimerMapper is cross-platform, freely available, and easily integrates with existing workflows. It offers a new perspective on the dynamic nature of predicted protein complexes, supporting researchers in the exploration of functional mechanisms and assembly paths.

## Introduction

Progress in artificial intelligence (AI) methods applied to biology and biotechnology has achieved significant milestones. AlphaFold2 (AF2) has increased protein structure prediction accuracy to unprecedented levels, providing the opportunity to obtain reliable structural information without the lengthy processes of experimental determination through NMR, X-ray crystallography, or Cryo-EM [1]. AlphaFold2-multimer (AF2m) extended this capability to protein complexes and protein-protein interaction prediction [2, 3]. Later, the implementation on cloud computing platforms has democratized their use in the scientific community, enabling researchers to run these models using only protein sequences as input [4]. Built on similar principles, other powerful models have emerged, including RoseTTAFold (RF), RoseTTAFold2 (RF2), and RoseTTAFoldNA (RFNA) [5–7]. The latest release of AlphaFold, AlphaFold3 (AF3), continues this progression by refining prediction accuracy and extending coverage to other molecules beyond polypeptides [8]. Each one of these advances aim toward greater versatility in modeling diverse atomic systems while improving computational efficiency and accuracy.

Despite these gains, there are still challenges that need to be addressed. For example, current AI models by themselves still fall short in capturing the dynamic nature of protein interactions and conformational diversity [9]. Furthermore, predictions of small complexes that involve just a few models are easily tractable and interpretable, but the systematic analysis and interpretation of structural ensembles for large unknown complexes remains difficult [10–12], particularly when the only information of a system is a list proteins sequences. Complexity increases further when transient or dynamic interactions and conformational changes are considered, as these can cause proteins to associate or dissociate from their partners in a time and contextual dependent manner [13–15]. Additionally, competitive binding scenarios where different proteins interact with the same surface on other proteins create situations where simultaneous interaction within the same complex becomes physically impossible [16]. These factors can lead to ambiguity in model interpretation, making it difficult to extract biologically relevant stoichiometries or interaction patterns. Lastly, while some methodological approaches were developed [17, 18], another difficult to solve problem is the prediction of stoichiometric coefficients when prior experimental data or templates are unavailable.

A relevant approach is the combinatorial assembly implemented by CombFold, which addresses some of these issues by assembling large complexes through overlapping subcomplex predictions [10]. However, this method often requires some prior knowledge of complex stoichiometry and domain boundaries, ultimately generating static models that does not give an explicit interpretation of interactions dynamics and conformational variability [10, 19]. Nonetheless, we hypothesized that these overlapping structural datasets can also be used to explore how predicted interactions and conformations change depending on the modeling context, a process that we refer to as “context-dependent structure prediction analysis”, which can be applied as a complementary approach to obtain meaningful biological information. Hence, we developed a method and software to systematically sample the stoichiometric space, capture and interpret these variations.

Before the initiation of this work, we intended to predict the structure of unknown trypanosomatid nuclear complexes, with the aim of design wet-lab experiments [20]. Soon, we faced the challenges already mentioned (large assembly complexity, transient/dynamic interactions, conformational changes and stoichiometric coefficient inference), inspiring the development of MultimerMapper, a complementary but comprehensive approach to analyze structural variability in predicted protein complexes under different modeling conditions, with a particular focus on protein interfaces. We kept trypanosomatids as our testing background due to their significance as disease-causing organisms, our expertise in their complexes and their high evolutionary divergence from model organisms [21]. In particular, this divergence is reflected in the low protein sequence identity compared to model organisms, but conserving the structural features of core eukaryotic complexes [20], providing an excellent opportunity to test our approach’s robustness. With this methodology, we attempt to address common limitations in the field, especially for highly dynamic complexes with unknown stoichiometries [10].

## Results

### 1. The Concept of Predicted Dynamic Interactions

To understanding the principles of our approach, we will first define some ideas. MultimerMapper introduces the novel concept of “predicted dynamic interactions”, based on the hypothesis that protein-protein interactions (PPIs) can engage or disengage depending on the context in which they are modeled. The basis of this idea is to compare pairwise interactions across different prediction scenarios, *e*.*g*., with or without specific protein partners. If applied properly, this allows the extraction of valuable information about complex assembly dynamics and stoichiometry, minimizing as much as possible the reliance on prior knowledge about the system.

#### 1.1. The Predicted Dynamic Interactions Hypothesis

PPI patterns from simple dimeric predictions (2-mers) are compared with those in higher-order oligomeric predictions (N-mers) using graph theory. To illustrate this concept, consider a hypothetical system of three proteins: A, B, and C (**Fig. *1*a**). For the moment, assume that we can determine with certainty that an interaction between two of these proteins exists in a structural model, involving two specific surfaces (green checkmarks in **Fig. *1***). The actual method used by MultimerMapper to predict interactions based on the data generated by AlphaFold (AF) is detailed below (see section 4. Protein-Protein Interaction Detector), along with the approach used to detect the involved interaction surfaces (see section 6. Contacts, Multivalency & Clustering).

**Fig. 1.**
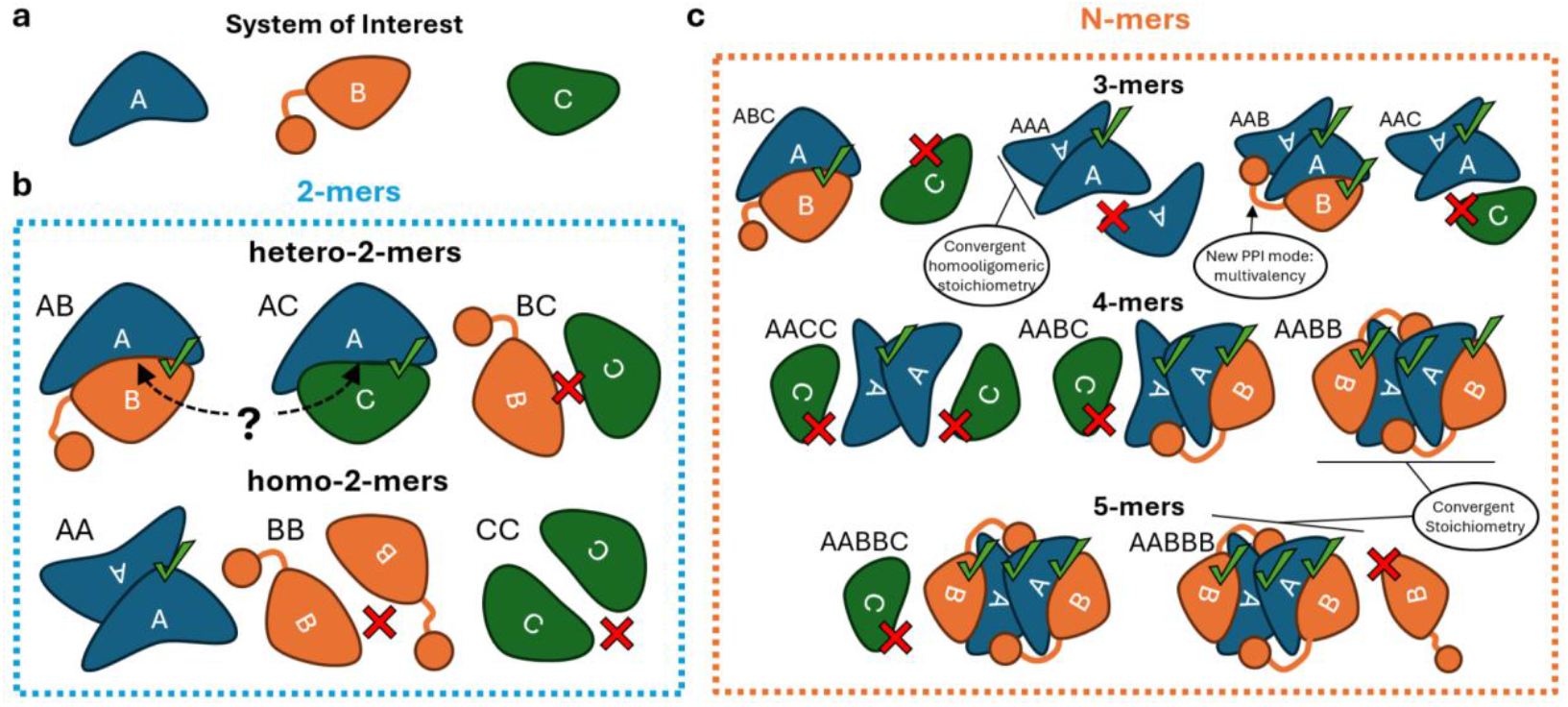
Hypothetical system with 3 proteins. (**a**) Monomeric structures of proteins A, B and C. B has two domains connected by a disordered region. (**b**) All possible dimeric predictions of the system. The presence or absence of interactions between proteins is highlighted with a green checkmark or a red cross, respectively. A co-occupied surface on A is indicated with arrows and a question mark. (**c**) Some possible oligomer (N-mers) structure predictions and the outcome of the interaction between subunits show with checkmarks and crosses. Key concepts described in this work that arise from the structural analysis of multimeric predictions are highlighted inside circles. These concepts include multivalency, convergence and convergent homooligomeric/multivalent states.

When all possible heterodimeric predictions are generated, we observe that A can interact with both B and C, while B and C do not interact with each other (**hetero-2-mers** in **Fig. *1*b**). Notice that the interaction surface of A in AB and AC is the same. One could ask: Could both interactions occur simultaneously? We can analyze their surfaces, align both complexes to check if there is any steric hindrance, etc., but we will take the following approach: predict the heterotrimer ABC and extract the PPI patterns in this new model (ABC **3-mer** in **Fig. *1*c**). There, only the A-B interaction persists, while the A-C interaction “disappears” with respect to the 2-mers. This simple comparison allow us to classify the A-B interaction as “static” (it is present in both 2-mer and 3-mer predictions) and the A-C interaction as “dynamic” (present in 2-mer but absent in 3-mer predictions). In this case, the dynamic behavior of the PPI reflects a steric hindrance, *i*.*e*., B and C might be competing for the same binding surface.

To systematically detect and analyze these dynamic PPIs, we developed a computational framework based on graph theory [22], with a set of rules to classify interactions and proteins based on their behavior (**Fig. 2a**). By comparing the graphs generated from 2-mers and N-mers predictions, we can classify PPIs and proteins as static or dynamic, along with the dynamic type (positive or negative). In our example, as the A-C interaction was present in the 2-mers, but disappeared in the computed N-mer, we classify its PPI as dynamic “negative”. This can happen when steric hindrance or conformational changes in higher-order complexes prevent certain interactions that were possible in simpler dimeric configurations [23]. In contrast, a dynamic “positive” PPI will happen when the opposite is true: not present in 2-mers but appears in N-mers. This can occur when the presence of additional proteins in a system creates new binding interfaces or induces conformational changes that promote interactions not observed in simpler dimeric predictions [14].

**Fig. 2:**
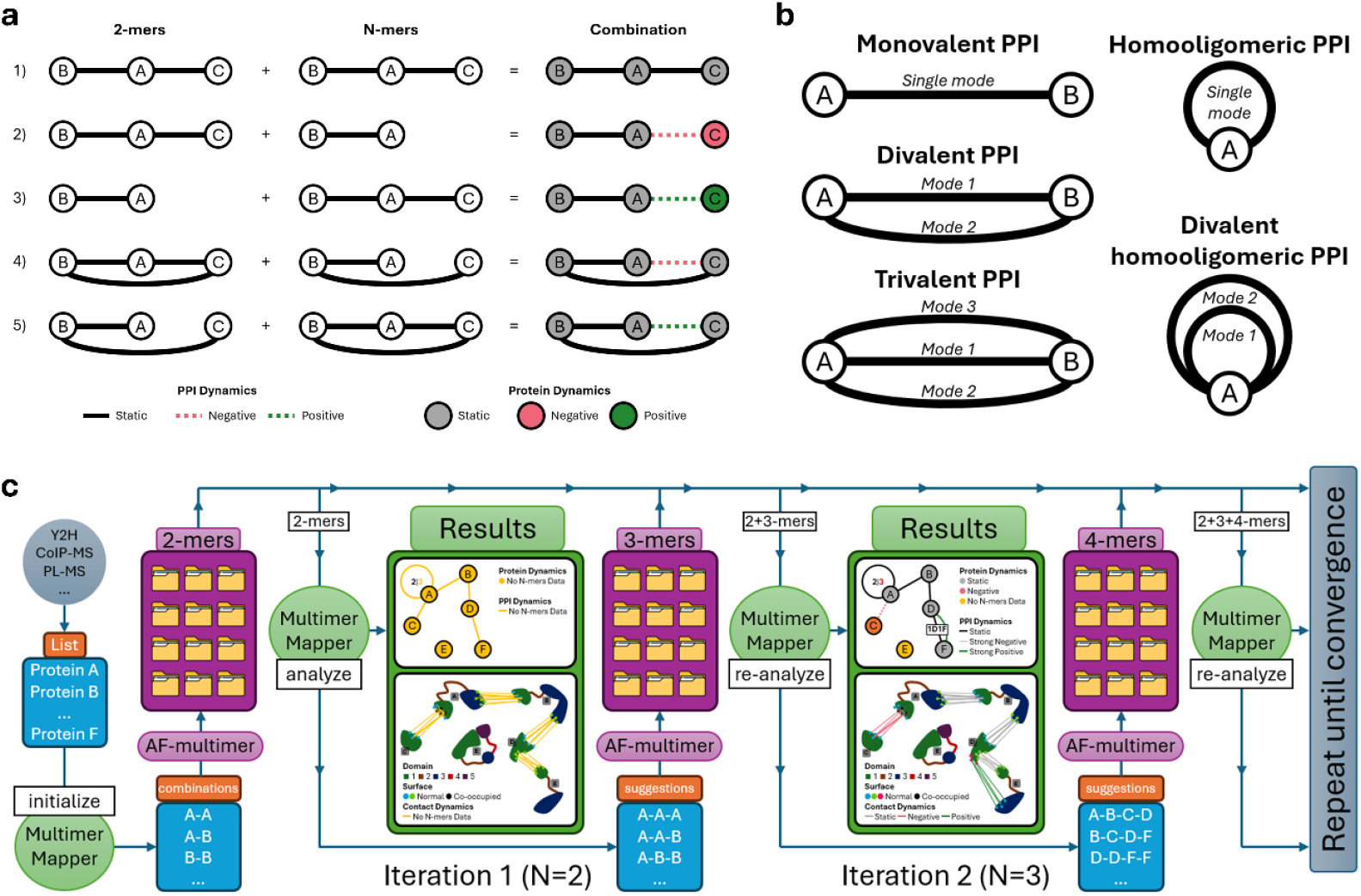
MultimerMapper’s PPI classification system and pipeline. (**a**) Logic of PPI and protein dynamics classification in a three proteins system. 1) Both interactions are present in both graphs. The proteins and interactions are static. 2) Both interactions are present in the 2-mer graph, but only the A-B edge is present in the N-mer graph. Therefore, C is a negative dynamic protein, and its interaction dynamics with B is negative. 3) The A-B edge is present in both graphs, and the B-C edge appears only in the N-mer graph. C is a positive dynamic protein, and its interaction dynamics with B is positive. This is a case of interaction activation induced by the presence of A. 4-5) Same case as 2 and 3, respectively, but involving a third interaction, A-C. In these cases, C is present in both combined graphs because it participates in a PPI that keeps this node in the final graphs. This makes protein C static, but not its interaction with B. (**b**) Graph representation of homooligomeric and multivalent interactions and their combinations. (**c**) MultimerMapper pipeline flow chart. A list of proteins with high probability of form a complex (*e*.*g*., coming from yeast-two hybrid, Co-Immunoprecipitation or Proximity Labeling experiments) are the initial input for MultimerMapper (initialize). All possible 2-mer combinations are generated (blue box) and the user must predict them using AlphaFold (2-mers purple box). Resulting prediction are processed with MultimerMapper (analyze). It will generate results and a new list of protein combinations (suggestions) that the user must predict (3-mers). Dimers and trimers are provided to MultimerMapper in a second iteration (N=3), producing more results and new suggestions. The process is repeated until the convergence of the system.

With this approach, AF applies its physicochemical/co-evolutionary “knowledge” to decide which PPI prevails (if any) in the combinations, while we explore the outcome and map their behavior on a combined graph. However, when multiple N-mers models contain the same protein under many different conditions, the possibilities are no longer binary, as the same interaction can appear in one context and disappear in another. Thus, the actual rules consider these possibilities as a PPI dynamics spectrum, based on the frequency of the PPI detected in the N-mers (see section 8. PPI Graph Converter).

#### 1.2. Incorporation of Homo-oligomerization and Multivalent Interactions

The framework also accounts for homo-oligomerization and multivalent interactions (more than one interaction surface between two proteins), two critical factors in determining complex stoichiometry [24, 25]. Homo-interactions are represented with self-connecting edges and multivalent interactions are represented as distinct paths between pairs of proteins (**Fig. 2b**). These two can also occur in combination and can be classified by their dynamic behavior independently. The method to detect multiple interaction surfaces based on multivalency is described below (see section 6. Contacts, Multivalency & Clustering).

Getting back to our thought example, after predicting the homodimers (AA, BB and CC) we only detect homooligomerization for A (**homo-2-mers** in **Fig. *1*b**). So, we can incorporate this information in a 2-mers PPI graph as an edge that self-connects A. Given that A was the only protein that interacted with itself and we don’t know anything about the system, we can explore the behavior for this homo-interaction in different trimeric contexts. For this, we generate combinations using two subunits of A and vary the third subunit. We predict the homotrimer AAA to see if it assembles into a stable trimer or if it has any effect in the homointeraction (AAA **3-mer** in **Fig. *1*c**). In this case, we see that the self-interaction still occurs, but one of the subunits is leaved aside. In other words, the evidence suggests that the homotrimer is unlikely to be formed, while adding an extra subunit does not disrupt the PPI. We can also predict the heterotrimers (AAB and AAC **3-mers** in **Fig. *1*c**). Interestingly, we notice that C does not interact with the homo-dimeric form of A in the AAC context. This suggests that something is happening with A when it engages interactions with itself that is preventing its interaction with C, maybe a conformational change. Moreover, B still interacts with the same surface on A in the AAB context, but a new interaction surface appears with another subunit of A. This is engaged via a different domain of B and can be added to the graph as a different edge.

This information, combined with the dynamic interaction data, provide insights into a potential regulatory role of protein C in the formation of an A-B complex. We will see that MultimerMapper is also capable of detecting conformational changes associated with distinct proteins when they are introduced in the predictions (see sections 5. Coordinate Analyzer and 8. PPI Graph Converter).

#### 1.3. Iterative Stoichiometric Space Exploration and the Convergence Hypothesis

Based in these ideas, MultimerMapper suggest combinations that the user must predict to explore the stoichiometric space in an iterative manner. The system is initialized with a list of proteins, generating all the possible 2-mer combinations that the user must predict (initialization step in **Fig. 2c**). Once provided, in the first iteration, the software determines which 2-mers engage PPI and combines the interacting pairs to suggest combinations of three proteins (3-mers). Once predicted, these 3-mers are then given back to MultimerMapper, which statistically identifies which are likely to form stable complexes. For now, let’s consider instability as having at least one red cross (**Fig. *1***). The actual algorithm to detect stability is described below (see section 7. Stoichiometric Space Exploration & Stoichiometry Prediction). Stable 3-mers are then chosen to generate new combinations (4-mers) that the user should predict for the next iteration.

As different combinations are discarded due to instability and do not generate new suggestions, this strategy assumes that the number of new combinations will decrease gradually at each iteration, as we approach the combination of proteins that represent what we call the “convergent stoichiometries”. As we will see, these generally match the reported functional stoichiometries of the underlying complexes.

On the other hand, these suggestions, focused on combinations that are relevant, maximize the identification of PPI dynamics, homo-oligomerization, multivalency and conformational changes within the system. However, it’s important to note that MultimerMapper cannot automatically detect non-convergent geometries (such as linear polymers like tubulin). In practice, users must identify them during the iterations, guided by MultimerMapper’s suggestions and visualizations, and remove them from the prediction to prevent infinite loops.

Computationally, the number of required predictions (P) is fixed for 2-mers and depends on the number of distinct polypeptides within the system (n): n (n + 1)

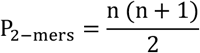

Meanwhile, P will vary for N-mers depending on the complexity of the system. In systems with no interacting pairs, N-mer predictions will be zero, while systems with potential for infinite growth (like linear polymers) could theoretically require an infinite number of predictions:

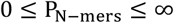

### 2. MultimerMapper Architecture

The software is flexible, accommodates diverse datasets, detects errors introduced by the user, and identifies gaps in data that are needed to estimate interaction dynamics and convergent stoichiometries. It provides output as both computational objects and user-friendly visualizations to help understand protein behavior and interfaces. The inner workings of MultimerMapper is illustrated in **Fig. 3**.

**Fig. 3:**
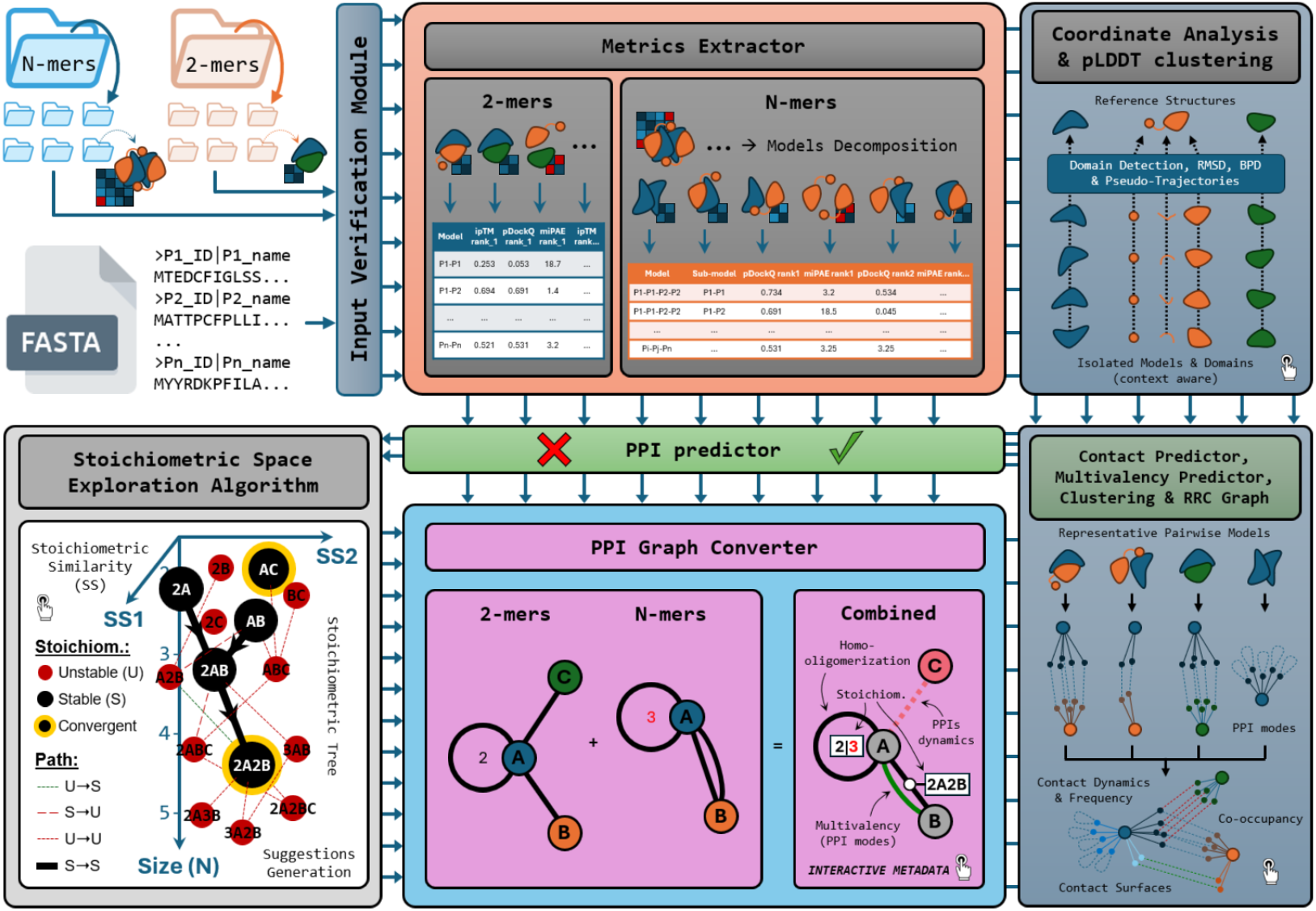
MultimerMapper main architecture and inner workings. Each core modules of the software and their inner workings are represented as boxes. A description of how each module works can be found in the main text.

The program’s minimum input requirements are:

1. A FASTA file containing amino acid sequences of the proteins present in the predictions. Each sequence header must include unique protein identifiers and a unique name or symbol for tracking and representations. For the initialization execution, this is the only input needed to generate the initial 2-mer combinations (**Fig. 2c**).
2. A directory containing all 2-mer predictions (both homo- and hetero-dimeric).
3. Optionally, a directory of N-mer predictions, which enables the detection of dynamic interactions and convergent stoichiometries.

The program is organized in modules as shown in **Fig. 3**. This setup supports an iterative pipeline where users initialize the system with MultimerMapper, generate the suggested 2-mers dataset and then run MultimerMapper again to detect interacting protein pairs in a first iteration. If the objective is simply to detect PPIs between 2-mers, this step will be enough. However, users can predict the output suggestion combinations and re-run MultimerMapper with this additional information (**Fig. 2c**). MultimerMapper will then identify dynamic interactions and notifies users if additional information is needed. This process is repeated until convergence is reached (*i*.*e*., no more suggestions).

It is worth noticing that these modules do not strictly represent the software’s architecture but closely resemble the actual code organization. In the following sections, a description of each core module inner-workings is given, along with the testing procedure and results for optimizing the parameters used.

### 3. Metric Extractor

After a verification step that identifies errors or inconsistencies (**Input Verification Module, Fig. 3**), a first module pre-process 2-mers and N-mers separately (**Metrics Extractor, Fig. 3**):

1. For 2-mers, it directly extracts all model metrics (pLDDT, pTM, ipTM, and PAE matrix). Key metrics such as miPAE (minimum interaction PAE, see section 4. Protein-Protein Interaction Detector) are computed, along with additional metrics that form part of MultimerMapper’s output (e.g., average pLDDT, pLDDT per protein). Values are obtained for each of five models predicted by AF.
2. For N-mers, each model is decomposed into its component pairs. For example, a 4-mer composed of two subunits of A and two of B (A_1_A_2_B_1_B_2_) is decomposed into all possible dimeric combinations (A_1_A_2_, A_1_B_1_, …, B_1_B_2_). This allows the computation of metrics on each resulting sub-model and its associated sub-PAE matrix.

These metrics are then passed to the next modules to continue the analysis process.

### 4. Protein-Protein Interaction Detector

The PPI detector module (**PPI Detector, Fig. 3**) takes in the extracted metrics to determine which protein pairs of each prediction engage in PPIs. The validation process for this module is described below.

#### 4.1. Generation of Benchmark Dataset

Given that our model organisms does not contain any gold standards to benchmark PPI prediction like, for example, in yeasts [26], it was necessary to generate a benchmark dataset. This was accomplished through manual curation of the literature, retrieving pairs of trypanosomatid proteins with experimental evidence of direct interaction engaging. The result was 175 pairs of interacting proteins. The negative dataset was generated using the following approach. Lists of proteins belonging to different eukaryotic complexes also conserved in trypanosomatids were generated and labeled according to the complex name and subcellular compartment. Pairs of complexes with unrelated functions and separate subcellular compartments were defined, such that their components have a very low probability of having cross interacting pairs. A pair of complexes was randomly selected, and for each complex, a protein was randomly chosen to generate a non-interacting pair. The process was repeated until 200 pairs were reached, without duplicates. The resulting data can be found in **Supplementary Table 1**.

#### 4.2. Structural Prediction and Metric Extraction for Benchmark

Dimers from the PPI benchmark dataset were predicted using a local version of AF2m (see methods) and the AF3 server [8]. Five models were predicted for each pair, ranking the models based on ipTM values. Associated metrics were extracted (pLDDT, pTM, ipTM, PAE) and two additional interaction metrics were computed: pDockQ, widely used and validated for interaction prediction [3]; and miPAE, which simplifies the PAE matrix by focusing on interaction regions. Now calculated by default by the AF3 server [8], miPAE identifies the minimum value in the PAE matrix for residue pairs across different chain pairs. The resulting data can be found in **Supplementary Table 2**.

#### 4.3. Optimization of Cutoff Values

To identify a cutoff values that maximize PPI prediction sensitivity and minimize non-specificity, we performed a ROC (Receiver Operating Characteristic) analysis using the interaction metrics (pDockQ, ipTM and miPAE). For each metric, we tested the effect of the number of models (N_models_) required to exceed the variable metric cutoff to consider an interaction. The resulting ROC plot can be seen in **Supplementary Figure 1**. We then extracted the cutoffs at false positive rates (FPR) of 0.01, 0.02, 0.05 and 0.1 and measured its sensitivity (**Fig. 4** and **Supplementary Table 3**).

**Fig. 4:**
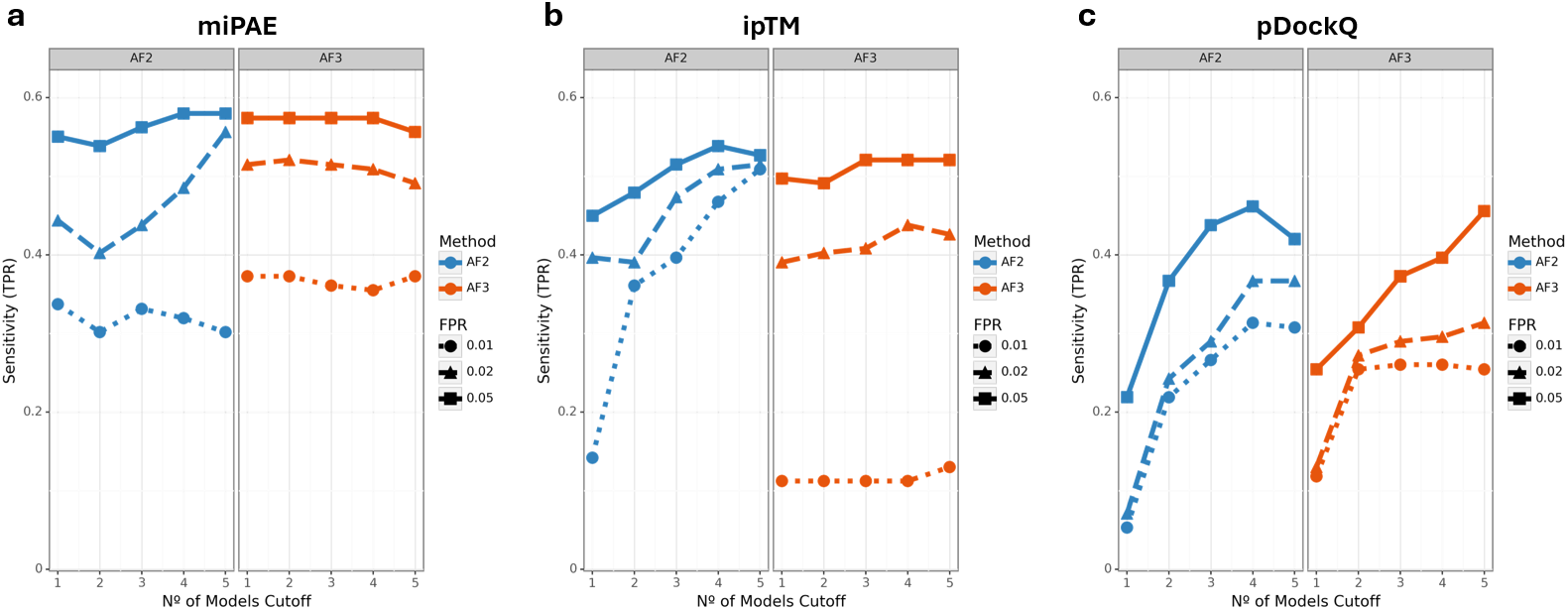
PPI prediction sensitivity profiles for different cutoff combinations and methods. Measured sensitivity using AlphaFold2-multimer (AF2) and AlphaFold3 (AF3) at different FPR levels (0.01, 0.02 and 0.05), N_models_ cutoffs and metrics. Profiles using the interaction metrics miPAE (**a**), ipTM (**b**) and pDockQ (**c**).

We observed that the best metric to predict PPIs is the miPAE for both methods (**Fig. 4a**), as it derives in the highest sensitivities, followed by the ipTM (**Fig. 4b**) and then the pDockQ (**Fig. 4c**). The same conclusion can be obtained by analyzing the AUCs (area under the curve) of the ROC plots (**Supplementary Figure 2**), being higher for miPAE. At FPR=0.05, AF2m and AF3 seems to have similar sensitivity profiles, but they varied in the obtained miPAE cutoffs (**Supplementary Table 3**).

By default, we have set the cutoffs of PAE ≤ 10 Å for AF2 and PAE ≤ 8.3 Å for AF3, with N_models_ = 4, which resulted in approximately 58% sensitivity at FPR=0.05 for both methods (**Fig. 4a**). However, these values can be modified by the user using a configuration file. During MultimerMapper executions, this criterion is applied directly for 2-mers. For N-mers, instead, it is applied to identify, at high level, if a PPI exists between any chain of both protein entities. For example, if a complex of 6 chains contain 2 subunits of protein A and 1 subunit of protein B, for any given model/rank, we consider that it has surpassed the cutoff if any miPAE of the two possible decomposed pairs for A-B surpass the cutoff. Then, at a later stage, within the execution, MultimerMapper identifies which of these chains are in physical contact (see section 6. Contacts, Multivalency & Clustering). In other words, at this step we only identify that a PPI between two protein entities exist within an N-mer combination, not between which chains.

### 5. Coordinate Analyzer

This module (**Coordinate Analyzer, Fig. 3**) processes the atomic coordinates, intramolecular PAE submatrices and pLDDTs of isolated chains from all models to evaluate how the protein conformation behaves depending on the modelling context. It first extracts a reference structure for each protein by selecting the overall chain with the best average pLDDT, which is used as reference point for several computations [10]. This module then:

1. Identifies well-structured domains and disordered segments by clustering the PAE matrix of the reference structure [27]. This can be done in two ways: automatic and semi-automatic. If user decide to use their own segments, a manual mode using a TSV file is available. It also identifies functional domains using InterProScan [28].
2. Calculates the Root Mean Square Deviation (RMSD) of isolated chains against the whole protein reference and isolated segments against a segment reference. Then computes full-length and per-segment pseudo-trajectories by sorting the models by increasing RMSD, using as reference the highest pLDDT model/segment.
3. Several metrics and visualizations are generated to understand how and why trajectories evolve.
4. Clusters the chains by their pLDDT profiles and analyzes how enriched are the different protein entities on each cluster. This allows associating pLDDT distribution patterns with the presence of specific proteins.

All computations are aware of the modelling context that produced each isolated chain, ultimately helping to associate variations in the computed metrics and protein conformation to the presence/absence of specific proteins. The information generated in this stage is then processed by several interconnected modules downstream.

#### 5.1. Domain Detection

The coordinate analyzer incorporates a domain detection algorithm that automatically identifies structural domains. This feature utilizes the intramolecular PAE matrix to partition proteins into regions of high internal structural coherence, offering flexibility through automatic, semi-automatic, and manual modes, allowing users to easily get fine-tune results. The software generates interactive 2D PAE matrix plots and 3D protein backbone representations in real-time, color-coded by detected domains and pLDDT, aiding in the interpretation and validation of domain assignments (**Supplementary Figure 3**).

In semi-automatic mode, users can interactively adjust a clustering resolution parameter (default 0.075) to optimize domain definitions for each protein, which are then stored in a resolution preset for future iterations. The algorithm incorporates a loop removal step, ensuring that small, potentially flexible regions are appropriately incorporated into larger domains, while larger loops are considered as unstructured domain boundaries. Lower resolution values will split proteins into fewer domains, and vice versa (**Supplementary Figure 3**).

It’s worth noting that detected structural domains in this context are not necessarily equivalent to the functional domain definition [29]. Rather, they represent continuous segments of proteins that exhibit similar behavior in the intramolecular PAE matrix of the reference structure. This approach allows for the separation of proteins into compact, well-structured defined segments and more flexible or disordered regions. This provides a rational framework to explore conformational changes and metrics variation for these segments instead of just whole proteins, increasing the detail of the posterior analysis. For functional domain identification, the InterProScan REST API is automatically queried using the protein sequences [28].

#### 5.2. RMSD calculations and Trajectories

MultimerMapper provides a comprehensive analysis of structural variation across different modeling contexts by computing RMSD values between each model chain and the reference structure. It tracks chain IDs and partners present in the models, offering insights into potential conformational changes associated with the modeling conditions.

AlphaFold has demonstrated the ability to predict conformational changes when the input MSA is sampled [30]. Following a similar rationale, our methodology generates different MSAs for each combination, but the sampling occurs at the multimeric level. Hence, to explore conformational heterogeneity as product of the contextual sampling, MultimerMapper constructs pseudo-trajectories by sorting monomeric models in ascending order of RMSD values relative to the reference structure (**Supplementary Video 1**). This is performed for both entire proteins and individual domains, providing a rich dataset for understanding structural variability and flexibility. When integrated with metadata such as radius of gyration (ROG) and pLDDT scores (see section 5.3), they offer deeper insights into how proteins respond to different modeling contexts.

#### 5.3. Bias in Partner Distribution (BPD) for RMSD trajectories and other metrics

To understand how and why these trajectories evolve, MultimerMapper builds several per frame features that can be interpreted along their corresponding trajectory structures. These features are provided as interactive plots (**Fig. 5a-f**) in parallel with an integrated trajectory visualizer to reproduce the pseudo-trajectory. This setup facilitates the exploration of the structures at each frame along the metrics features without relying on external software (**Supplementary Figure 4**).

**Fig. 5:**
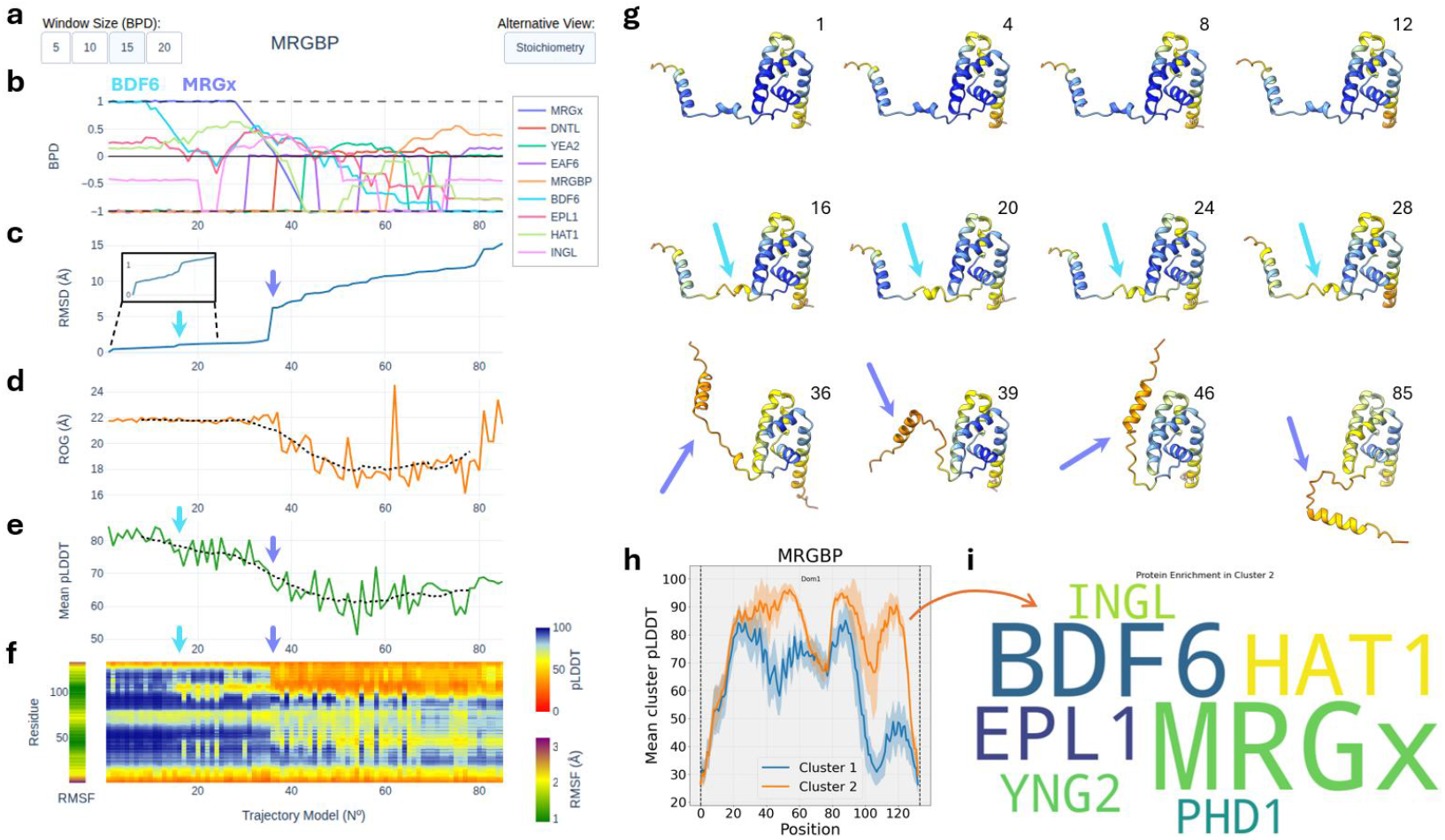
Data and visualizations generated by the coordinate analyzer module. (**a**) Interactive visualization with detected domains for the test protein BDF5 [20]. The default resolution value accurately detects each domain and disordered segments. (**b**) Diminishing the resolution value results in fewer domains detected, and vice versa. (**c**) Example RMSF curve for MRGBP of *T. brucei*. (**d**) Values of RMSD, ROG and mean pLDDT along the pseudo-trajectory of MRGBP. The green arrow indicates the point at which a very small change in the metrics occurs, while the blue arrow indicates a sudden spike in RMSD, associated with a drop in ROG and mean pLDDT. (**e**) pLDDT heatmap for the trajectory of MRGBP. The same green and blue arrows from panel d indicates the respective points of changes in per-residue pLDDT. Trajectory positions after the blue arrow show low pLDDT values for the C-terminal residues. (**f**) Bias in Partner Distribution of MRGBP pseudo-trajectory. Representative trajectory models are shown at the bottom, colored by pLDDT. Initial trajectory models show high bias for the presence of BDF6 and MRGx. When BDF6 bias disappears, slight changes are seen in structural conformation. When MRGx bias disappears, radical changes in pLDDT and conformation occurs, particularly in the C-terminal α-helix connected by a loop, which starts to oscillate. (**g**) Models chains pLDDT clustering results for MRGBP. MultimerMapper detects two pLDDT profiles (cluster 1 and 2) for MRGBP models. (**h**) Partner enrichment analysis of cluster 2 from panel g. The size of protein names represents their frequency in the cluster. MRGx is the most enriched partner (representative), followed by BDF6.

The RMSD, besides being used to generate the trajectory, provides a good measure of how much the conformation changes along the trajectory. Sudden changes in RMSD may indicate major conformational changes occurring at specific points (**Fig. 5c**). Another informative metric is the Radius of Gyration (ROG), which assess the overall compactness of the structure throughout the pseudo-trajectory [31]. Variations in ROG may indicate transitions between more compact and extended conformations, reflecting potential structural rearrangements or binding events (**Fig. 5d**). Mean pLDDT provides insights into average changes in overall local confidence [1]. Abrupt changes in mean pLDDT may indicate the engagement/detachment of an interaction at specific points that affect confidence, whereas stable values can suggest structural stability along the whole trajectory (**Fig. 5e**). Additionally, a per-residue pLDDT heatmap for the trajectory is provided to pinpoint specific regions responsible for confidence variations (**Fig. 5f**).

To correlate these structural and metrics variations through the trajectory with the presence/absence of specific proteins, we developed a novel measure that we call Bias in Partners Distribution (BPD). This metric evaluates whether certain proteins are preferentially enriched or depleted in specific frames of the pseudo-trajectory. BPD is computed by creating sliding windows of trajectory frames and analyzing the frequency of each protein in the window relative to its overall frequency (bias). A bias score of 1 indicates that a protein is strongly enriched in a given window, while a score of −1 means it is completely absent. A value around 0 suggests a random distribution, meaning the presence of the protein is not significantly biased within the trajectory window (**Fig. 5b**). This measure helps identify whether specific interaction partners are associated with particular conformational states, providing insights into potential binding preferences, induced fit mechanisms, allosteric regulation or assembly paths.

The Root Mean Square Fluctuation (RMSF) gives measure of how much each residue fluctuates throughout the trajectory [32], revealing regions with the highest conformational flexibility across all predictions chains. For the interactive visualizations, RMSF is provided as a per-residue heatmap (**Fig. 5f**, left).

For example, the protein from **Fig. 5a-f** is MRGBP from *Trypanosoma brucei*, analyzed within the context of the entire NuA4 complex [20, 33]. Previous molecular dynamics simulations and experimental data demonstrated that MRGBP binds sequentially to the proteins MRGx and BDF6 to form the conserved eukaryotic subcomplex TINTIN. MRGBP is too unstable to interact with BDF6 in its free form; instead, it first binds to MRGx, which stabilizes its C-terminal segment and enables its subsequent interaction with BDF6 [20]. This same sequential binding pattern emerges in our pseudo-trajectory analysis, where BPD scores indicate a high bias for both MRGx and BDF6 in early frames, followed by the sequential decrease of BDF6 BPD values, and then MRGx (**Fig. 5b**). The initial depletion of BDF6 (frame at which the BDF6 gets a BPD of zero) is associated with small increases in RMSD and small decreases of pLDDT (light blue arrows in **Fig. 5c-g**). At the conformational level, the change occurs in the first section of the extended segment of MRGBP (light blue arrows in **Fig. 5g**). After the depletion of MRGx, an abrupt change in conformation, reduction of the ROG and drop in confidence scores can be observed (purple-blue arrows). These mirrors previous molecular dynamics and experimental observations [20], now captured through a modeling-based approach.

Another useful visualization to identify trends in protein context associated with specific conformations shows how the stoichiometric coefficients of each protein are distributed along the trajectory using windows. This is complementary to the BPD and gives information about the specific amount of each protein in the frames of the pseudo-trajectory. The interactive version has buttons to change between the Stoichiometry and BPD visualization using windows (**Fig. 5a**).

#### 5.4. pLDDT clustering

Complementary, a pLDDT clustering analysis step groups the models chains based on their local confidence scores, revealing patterns in model confidence across different contexts (**Fig. 5g**). Similarly to the BPD, this is complemented by a “partner enrichment analysis”, which identifies specific partners associated with specific confidence profile clusters. The analysis computes the frequency of co-occurrence of proteins within each pLDDT cluster, detecting overrepresented partners.

To visualize these patterns, the module generates word clouds for each cluster (**Fig. 5h**), highlighting the most common partners associated with different confidence profiles. This makes it easier to identify which partners frequently appear in models with similar pLDDT patterns. Additionally, assigning a representative partner to each model simplifies interpretation, enabling a quick assessment of the dominant interaction context for each structural prediction. By integrating these analyses, the approach uncovers potential relationships between protein-protein interactions and model confidence, offering valuable insights into how interaction contexts influence local model quality. For example, in **Fig. 5h-i**, similarly to what was observed in **Fig. 5b-g**, we can see that MRGx and BDF6 are the most enriched proteins in the orange cluster, followed by HAT1 and EPL1. In other words, when MRGx and BDF6 is present in the models that contain MRGBP, MRGBP has better pLDDT values.

### 6. Contacts, Multivalency & Clustering

Another module of MultimerMapper (**Contact predictor, Multivalency Predictor, Clustering and RRC graph, Fig. 3**) is in charge of analyzing the interaction surfaces between protein pairs. It extracts the interchain residue-residue contacts (RRC) of each possible pairwise sub-model as contact matrixes, identifies multivalency, clusters the contact matrices into distinct PPI modes and generates RRC graph representations. Individual RRC pairs are also classified as dynamic and static, simultaneously tracking residues involved in multiple contacts that could potentially cause steric hindrances (surface co-occupation). Interactive representations to explore the results are also generated.

#### 6.1. Interprotein Residue-Residue Contacts Prediction

Interchain RRCs are predicted on both 2-mer and decomposed N-mer pairs that were detected engaging PPIs in their corresponding models (see section 4. Protein-Protein Interaction Detector). RRCs are considered null for protein pairs on models that PPI detection resulted negative. The method combines information from the interchain PAE matrix (iPAE) and the interchain residue distances (distogram) to identify residue pairs that are in contact (**Supplementary Figure 5**). It consider a residue pair in contact if the error is lower than the PAE cutoff used for PPI detection and their centroid center of masses are sufficiently close (less than 8 Å).

#### 6.2. Multivalency Prediction

Once identified which chains are in contact inside each model, MultimerMapper predicts which protein pairs engage multivalent interactions by counting how many isolated chains are in contact with more than one of the corresponding opposite pair at the same time (**Supplementary Figure 6a**). To distinguish real from spurious multivalency, different metrics derived from the counts were built and tested using a benchmark dataset consisting of multivalent and monovalent protein pairs time (**Supplementary Table 4**). The best performant metric was the fraction of multivalent chains (FMC) of each pair (**Supplementary Figure 6b-c**), obtaining a sensitivity of 73% and a specificity of 93% using an FMC cutoff of 0.2.

#### 6.3. Contact Matrix Clustering

For multivalent protein pairs, MultimerMapper predicts the number of distinct PPI binding modes (see **Fig. *2*b**) by clustering their contact matrixes. For this, we developed several contact matrix clustering methods and tested them against the multivalent pairs of the benchmark dataset to compare their performance against random models. This benchmark dataset consisted of protein pairs for which the number of binding modes was curated manually. The performance of each method was based on the correct identification of binding modes numbers using recall, precision and accuracy (**Supplementary Figure 7**). The best method performs hierarchical clustering over a distance matrix built using RMSD distances of the *α*-carbons of contact involved residues between matrix pairs (see methods).

The objective of merging individual matrices into clusters by similarity it to obtain an average representation of the binding interface of each mode. It also allows computing how frequent each interaction mode is represented in the N-mers, and the frequency of individual contacts within each cluster. The implementation is also aware of which combinations of proteins contributed more to each cluster and, by comparing which context produced which contact mode (2-mer versus N-mers), it is possible to associate presence/absence of certain partners to certain PPI modes. Moreover, individual contacts can be classified into static, dynamic positive and dynamic negative.

Once the contact matrixes are clustered, the software identifies a representative structure as the pairwise model with the best contact matrix overlap to the average contact matrix of the cluster. For example, **Fig. *6*a-b** show the representative structures for each identified cluster of the pair SEC31-SEC31 and SEC13-SEC31, respectively, processed with MultimerMapper until convergence of the validation system composed of the protein entities SEC13 and SEC31. These two proteins form the outer coat of the core eukaryotic complex COPII, a cage like complex involved in the budding of vesicles that is essential for secretory pathways [34]. The identified clusters reflect the flexibility of SEC31 and heterogeneity of interactions, something that has been experimentally observed, which is key for the function of the cage [35]. SEC31-SEC31 Cluster 0 and 2 representatives are comparable to previously observed conformations of SEC31, while cluster 1 and 3 representatives seem to be transition conformations between cluster 0 and 2 (**Fig. *6*a**). Meanwhile, the clusters for SEC13-SEC31 show how SEC31 can bind SEC13 using only its C-terminal, only its N-terminal or with both simultaneously (**Fig. *6*b**). Contact cluster information can be analyzed in parallel with the pseudo-trajectory of SEC31, which captures this conformational transition in more detail (**Fig. *6*c and Supplementary Video 2**).

**Fig. 6:**
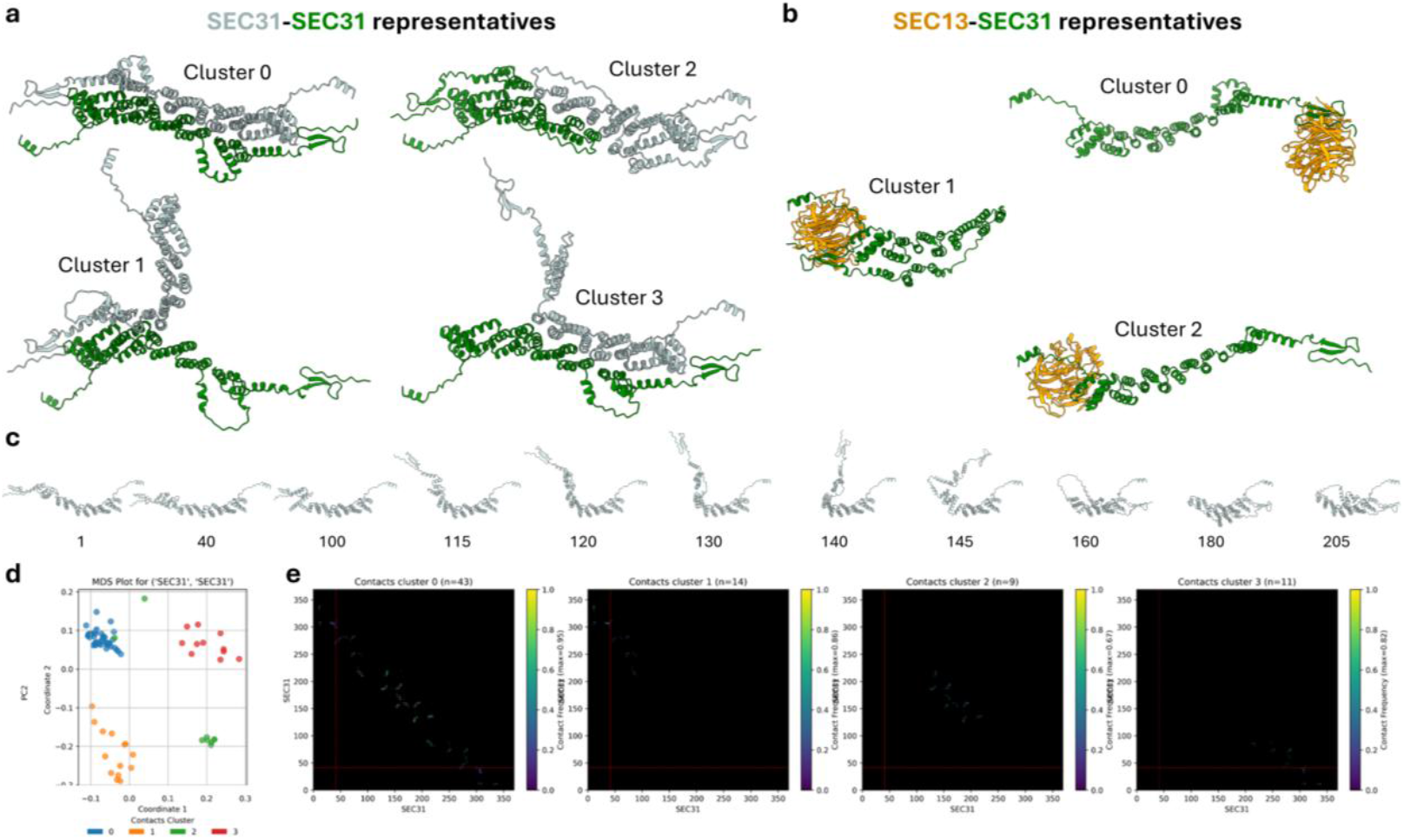
Contact matrix clustering. Example results of the contact matrix clustering algorithm for the validation system composed of the protein entities SEC13 and SEC31 from *Leptomonas seymouri*. (**a**) Clusters representative structures of the pair SEC31/SEC3. (**b**) Clusters representative structures of the pair SEC13/SEC31. (**c**) Pseudo-trajectory models showing the conformational trajectory of SEC31. The numbers below the structures are the corresponding frames on the pseudo-trajectory. (**d**) MDS plot colored by contact cluster for the pair SEC31/SEC31. Each marker represents a decomposed pairwise model. Coordinates 1 and 2 are low-dimensional embeddings that place models so that those closer have small RMSD and those far apart have large RMSD. (**e**) Average contact matrixes of the contact clusters from panel **a**. The size of each cluster is shown between brackets (n) and the frequency of each RRC in the cluster is color coded. The maximum observed contact frequency is also provided (max).

A multidimensional scaling (MDS) plot that integrates the RMSD distances to see relationships between pairwise models and the clustering results is also provided (**Fig. *6*d**), along the corresponding average contact matrixes of each cluster (**Fig. *6*e**). All these outputs are provided as interactive visualizations that the user can explore to understand the contextual behavior of the proteins.

For monovalent pairs, they are considered as interacting via a single binding mode, and their contact matrixes are grouped into a single cluster.

#### 6.4. Residue-Residue Contacts Graph Representation

To visualize what are the available contact surfaces between proteins, a 3D graph representation of the RRC clusters is generated and included as default output (**Supplementary Figure 8**). It represents each protein in 3D space using the reference structure, with individual RRCs depicted as lines that connect the residue centroids that were predicted to be in contact. The thickness of these lines is proportional to the frequency of the RRCs and are colored following the rules described in **Fig. *2*a** at the residue-level (static in gray, dynamic negative in red and dynamic positive in green). A force field algorithm is executed to orient the surfaces properly and avoid collisions. The visualization is interactive, with different color schemes (by pLDDT, by domain, by residue, etc.), protein styles (cartoon, spheres, etc.) and other customizable features available to explore the predicted RRCs between proteins.

### 7. Stoichiometric Space Exploration & Stoichiometry Prediction

A module of MultimerMapper (**Stoichiometric Space Exploration Algorithm, Fig. *3***) is in charge of suggesting combinations, finding the highest stable stoichiometries (convergent stoichiometries) and deciding when overall convergence has been reached. Besides predicting stoichiometries, this module is key to generate a rich dataset of models coming from different contexts that can be decomposed and analyzed by the rest of the software.

#### 7.1. The Stoichiometric Space Exploration Algorithm

**Fig. *7*a** shows a schematic representation of the algorithm for a system of two proteins, A and B. It starts with the initialization of the system and the prediction of all 2-mers. Once provided, MultimerMapper decides which 2-mers form stable complexes. It then use stable stoichiometries as base to construct new children stoichiometries that must be predicted for the next iteration. These are built by taking the combination of proteins in the stable parental stoichiometry and adding +1 of each protein entity to each new child. Meanwhile, unstable stoichiometries do not generate children. By default, only protein entities that are detected forming a fully connected PPI network with the proteins in the parental stoichiometry are used. For example, suppose the stable stoichiometry 3B has been detected in a system of 5 protein entities (A, B, C, D and E). A, B and C has been detected interacting with each other, while D and E interact in a different subnetwork. Then, by default, 3B will generate the children 1A3B (+A), 4B (+B) and 3B1C (+C). The software end up building a directed graph were stoichiometries grow by 1 unit at each iteration, exploring only informative combinations that have potential to be stable. Each stoichiometry can have multiple parents, and the biggest stable stoichiometries found at each branch that generates all unstable children is considered a convergent stoichiometry, which usually matches the reported stoichiometry of the system.

**Fig. 7:**
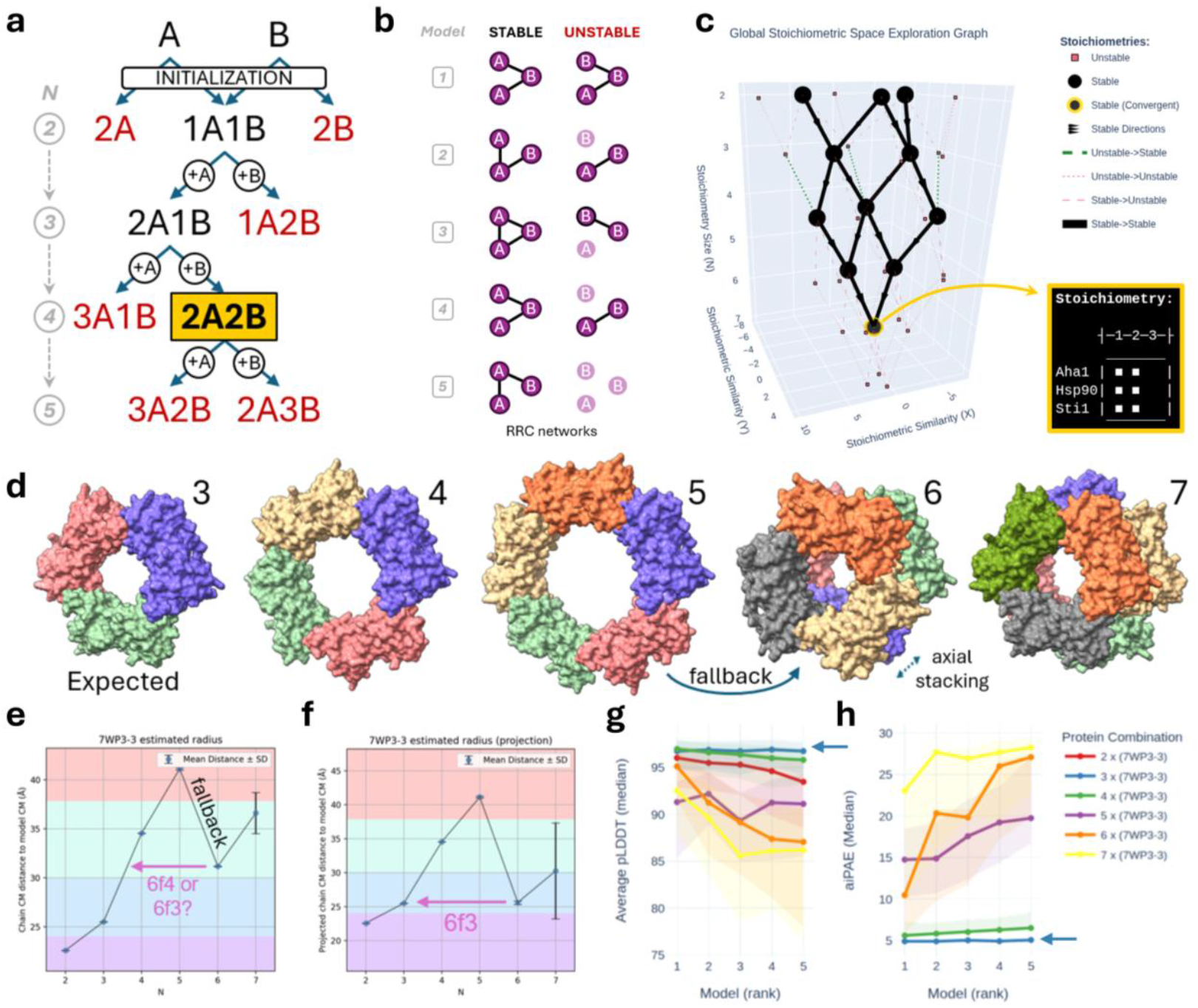
Stoichiometry prediction methods. (**a**) Schematic representation of the stoichiometric exploration algorithm for a simple system of 2 protein entities that converges in a 2A2B stoichiometry (gold). In the initialization, all 2-mers are computed. At the first iteration (N=2) MultimerMapper decides which pairs are stable. Unstable stoichiometries (red) do not generate children. Stable stoichiometries (black) generate children stoichiometries by adding +1 of each protein entity that has been detected forming a connected network with the proteins from the parental stoichiometry. These children stoichiometries (suggestions) must be predicted and included in the next iteration (N=3). The process is repeated until all the terminal stoichiometries from the hierarchies are unstable (convergence). (**b**) Definition of stability for an N-mer of 3 chains, A, B and C. For each model, a RRC network is built. Stable N-mers generate fully connected networks reproducibly (left). Unstable N-mers systematically leave chains outside the RRC network (right). (**c**) Interactive stoichiometric space exploration graph for the system composed of Hsp90, Aha1 and Sti1 from *T. brucei*. Each marker represents a stoichiometry (combination). Stable stoichiometries are represented as black circles and unstable as small pink squares. The convergent stoichiometry is represented with a golden outline and the interactive metadata with the stoichiometry is shown in the inset. Lines show the hierarchy between stoichiometries from different N values. (**d**) Rank 1 structures for test subject 7WP3. The expected stoichiometry structure is highlighted (N=3), along with the N value at which the symmetry fallback is detected. (**e**) Estimated radius of models shown in panel **d**. Bars represent the standard deviation (SD). (**f**) Estimated radius of models shown in panel **d** using 2D projection. (**g**) Average pLDDT confidence profile for test subject 7WP3. An arrow points to the expected stoichiometry confidence profile. The legend for each curve is shown in panel **h**. (**h**) Average interchain PAE (aiPAE) profile for test subject 7WP3.

2-mer stoichiometries are considered stable if they were predicted to be interacting (see section 4. Protein-Protein Interaction Detector). To define an N-mer stoichiometry as stable or unstable, MultimerMapper first builds a RRC network for each of the five models predicted by AlphaFold. Then, a stoichiometry is considered stable when there is, for multiple independent networks, at least one path connecting any pair of chains (**Fig. 7b**).

The algorithm was tested using different values of PAE and N_models_ cutoffs against a dataset of complexes with a wide range of known expected stoichiometries (monomers, homodimers, heterodimers, homotrimers, heterodimers, etc.). The PAE cutoff is the minimum interchain PAE needed to consider a residue pair in contact if they are less than 8 Å apart, and the N_models_ cutoffs is the minimum number of networks with at least one path connecting any pair of chains. We tested the algorithm using the PAE cutoffs obtained for the PPI prediction at 0.01, 0.02, 0.05 and 0.1 FPR, varying the N_models_ cutoff, to see which configuration identified the highest proportion of expected stoichiometries. Results can be seen in **Supplementary Figure 9**.

After every MultimerMapper run, an interactive 3D visualization showing the evolution of the stoichiometric space exploration is provided, allowing users to visually explore the results of the algorithm. For example, **Fig. 7c** shows the results for the system composed of the chaperone Hsp90 and the co-chaperones Aha1 and Sti1 from *Trypanosoma brucei* [36]. Hsp90 arranges into a homodimeric central complex that interacts with two subunits of Aha1 and two subunits of Sti1 throw distinct surfaces, forming a heterohexameric complex [37, 38], which is the convergent stoichiometry detected by MultimerMapper (**Fig. 7c**).

For the special case of systems were the involved complexes grow by addition of dimers and not just isolated monomers, a parameter was included to control the order of increments. The clearest example is the nucleosome, that physiologically assembles through the addition of histone dimers H2A:H2B and H3:H4 [39]. AlphaFold seems to replicate this behavior, as the nucleosome system converges in the isolated dimeric complexes if this parameter is set to add the default increments of 1 (**Supporting Figure 10a**). In contrast, MultimerMapper finds the expected convergent stoichiometry of the nucleosome (2:2:2:2) at N=8 when the parameter is set to work using increments of 2 (**Supporting Figure 10b**). This parameter works by generating children stoichiometries that not only adds +1 of each protein but also adds +2, using the stoichiometries of the stable 2-mers to each stable stoichiometry (see methods). It can be set to any integer, for example, to add up to +3 increments (adds stable 2-mers and 3-mers) or +4 (2, 3 and 4-mers), to cover other potential types of complex assemblies.

#### 7.2. Fallback Detection

By default, we included extra analysis based on some observations. Following the stoichiometric space exploration, for homooligomeric subsystems (just one protein entity), the software will systematically examine if the available homooligomeric predictions can form stable structures with increasing number of copies (homodimer, homotrimer, homotetramer, etc.). For some cyclical homooligomeric complexes, depending on the configuration, MultimerMapper detects convergent stoichiometries that are bigger than the expected size.

After carful exploration of their predicted structures, we noticed that AlphaFold predicts high confidence structures that extends beyond the expected stoichiometry, forming increasingly bigger cyclic structures. However, there is a point at which they experience a “fallback” to a more compact architecture with stack repetitions of the expected stoichiometry. For example, validation subject 7WP3 forms rings of increasing size with 3, 4 and 5 subunits, until it “falls back” at N=6, to a structure composed of two stacked elements of the structure predicted at N=3, which is the expected convergent stoichiometry (**Fig. *7*e**). Then, at N=7, one of the stacked substructures starts growing again. It seems that AlphaFold finds that it has room to add extra subunits beyond the expected stoichiometry, up to the fallback point, at which the structure cannot physically accommodate more and decides to split the complex into two. This same behavior was observed for the validation subject 7V4J, which is expected to form a cyclical pentamer, but grows beyond the pentamer and falls back to a stacked structure of 5 subunits (**Supplementary Fig. 11**).

Therefore, we built a fallback detection algorithm applied to homooligomeric proteins that can help to identify stoichiometries in certain situations. It consists of computing the estimated radius at each N value (see methods) and then detect if there is a significant drop in the successive radii (**Fig. *7*e**). If there is a significant drop, the algorithm tries to identify if the confidence interval of the falling back model’s estimated radius covers the mean estimated radius of any smaller N structure. As seen in **Fig. *7*e**, the interval does not cover any of the previous radius. However, by applying a 2D projection before estimating the radius at each N value (see methods), the estimated projected radius confidence interval covers perfectly the estimated projected radius at N=3 (**Fig. *7*f**). This information is added to the interactive 2D graph using the nomenclature YfZ (Y falls back to Z), where Y is the N value at which the fallbacks occurs and Z is the one to which Y falls back.

#### 7.3. Pairwise Confidence Profiles

Another observation made during the development of MultimerMapper was the tendency of average confidence metrics computed for protein pairs to peak at the expected stoichiometries. For example, **Fig. *7*g** shows the per-rank average pLDDT values of validation subject 7WP3 (**Fig. *7*d**) for different stoichiometric combinations. There, it can be seen that the average pLDDT profile gets maximized for the expected stoichiometric combination (N=3). Something similar happens for the average interaction PAE (aiPAE), being minimized at N=3 (blue arrow).

Due to this, we included these confidence profile plots by default to visually compare the average, standard deviation, median and interquartile ranges of the mean pLDDT, the miPAE and the aiPAE of pairwise interactions. These graphs helps to easily determine which combination of proteins generate the pairwise interactions with the best confidence metrics. In the case of systems or subsystems formed by homooligomeric or multivalent protein pairs, it can help to decide which is the most probable stoichiometry.

### 8. PPI Graph Converter

This module (**PPI Graph Converter, Fig. 3**) transforms direct interaction information between protein entities retrieved by the PPI and RRC predictors into graph representations, following the rules described in section 2 (**Fig. *2*a-b**). Additional statistical analysis are performed to explore the significance of associations between the presence or absence of certain partners with specific PPI patterns and conformational states.

#### 8.1. Direct Interaction Network Representation

MultimerMapper creates a network where each vertex is a protein entity and each edge is a particular PPI mode, both with distinct attributes. To build this graph, it first maps all the proteins that were involved in interactions, leaving outside of the network those that do not interact with any protein. Then, for each detected PPI mode between proteins, it analyzes if it was detected in the 2-mers, if it was detected in the N-mers and the relative frequency of the interaction in the N-mers. It also highlights if the PPI was not tested (no data), *i*.*e*., if the pair of proteins involved in the interaction were predicted together in at least one model, distinguishing this between 2-mers and N-mers. At the end, the module classifies each PPI using the **Table *1***.

**Table 1:**
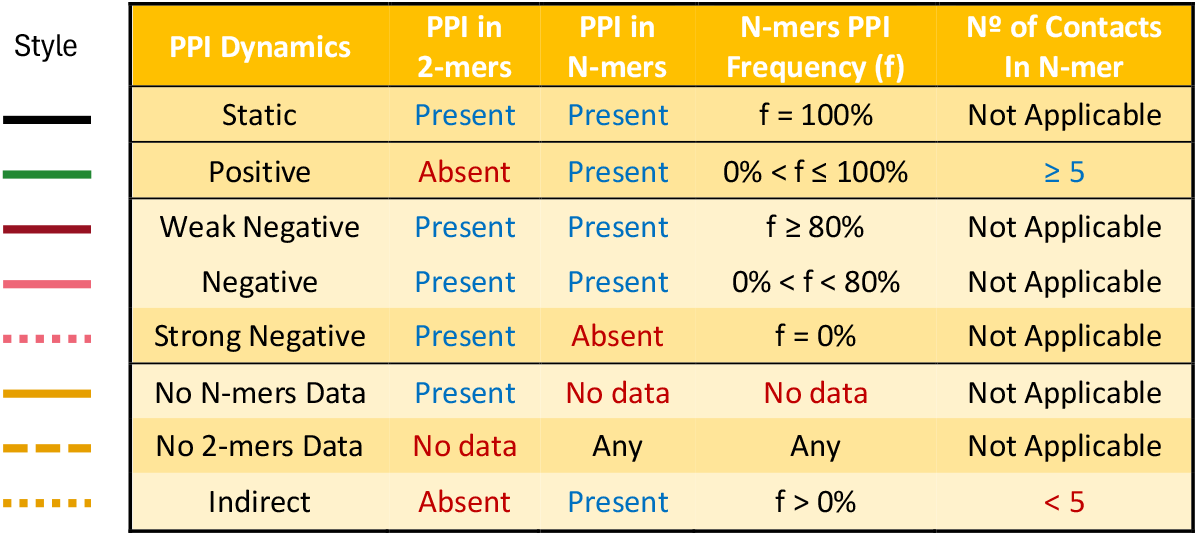
Extended PPI classification system. PPI Dynamics represents the classification name of the PPI. 2-mers and N-mers columns represents the binary outcome of both 2-mers and N-mers PPI analysis, respectively. N-mers Variation is the percentage of predictions that detected the PPI to be present in N-mers. N-mers Contacts is the minimum number of contacts needed between the PPI pair. The Style is the default representation in the interactive PPI graph produced by MultimerMapper. By default, Indirect interactions are removed from the combined graph.

| Style | PPI Dynamics | PPI in 2-mers | PPI in N-mers | N-mers PPI Frequency (f) | Nº of Contacts In N-mer |
| --- | --- | --- | --- | --- | --- |
|  | Static | Present | Present | f = 100% | Not Applicable |
|  | Positive | Absent | Present | 0% < f ≤ 100% | ≥ 5 |
|  | Weak Negative | Present | Present | f ≥ 80% | Not Applicable |
|  | Negative | Present | Present | 0% < f < 80% | Not Applicable |
|  | Strong Negative | Present | Absent | f = 0% | Not Applicable |
|  | No N-mers Data | Present | No data | No data | Not Applicable |
|  | No 2-mers Data | No data | Any | Any | Not Applicable |
|  | Indirect | Absent | Present | f > 0% | < 5 |

This assigns a different style to each PPI dynamic type, representing PPI frequency as edge width. In this way, it is possible to determine easily which are the most important or core interactions of the complexes, which interactions tend to appear (dynamic positive) or disappear (dynamic negative) in more complex scenarios and in which frequency. By default, MultimerMapper removes indirect interactions from the graphs. For protein dynamics, it only analyzes the overall presence or absence in the 2-mer and N-mer graph (**Fig. 2a**), not its relative frequency (see methods).

Everything is integrated into an interactive visualization that allows the user to explore useful metadata about the proteins and their interactions, like the exact N-mer frequency or the number of pairwise models that were detected interacting via particular PPI modes. For interactions with the potential to from higher order subassemblies (like homooligomers and multivalent interactions), the stability of the different combinations is shown in insets that can be turned on and off.

For example, **Fig. *8*a** shows the resulting PPI graph for the chaperone complex from **Fig. *7*c**. Hsp90 is known to form a dimeric subcomplex [36]. This is captured by MultimerMapper and represented by a self-connecting edge with an inset that indicates that the dimer is stable but the trimer is not. Meanwhile, the homooligomerization PPI is predicted to be present in all possible contexts. Hence, the edge is classified as static. Each Aha1 and Sti1 subunit interact with both subunits of the core Hsp90 dimer, resulting in a multivalent interaction. Only one of these binding modes is predicted by AlphaFold in Aha1-Hsp90 and Sti1-Hsp90 2-mers, while the other only appears in the N-mer combinations, represented as a green edge. Both are known to form heterotetrameric assemblies with two subunits of Hsp90 and two subunit<s of Aha1 or Sti1 [37, 38]. Similarly to what happens to the homooligomerization, this is also captured by MultimerMapper, where the highest stable stoichiometry is a tetramer, represented with the nomenclature 2P2Q. Notice that Aha1 interacts with itself, but the dimer and trimer are unstable, and the PPI edge is dynamic positive. This is due to the engagement of additional interactions that are only possible in the N-mers, when two Aha1 subunits interact simultaneously with the Hsp90 dimer.

**Fig. 8:**
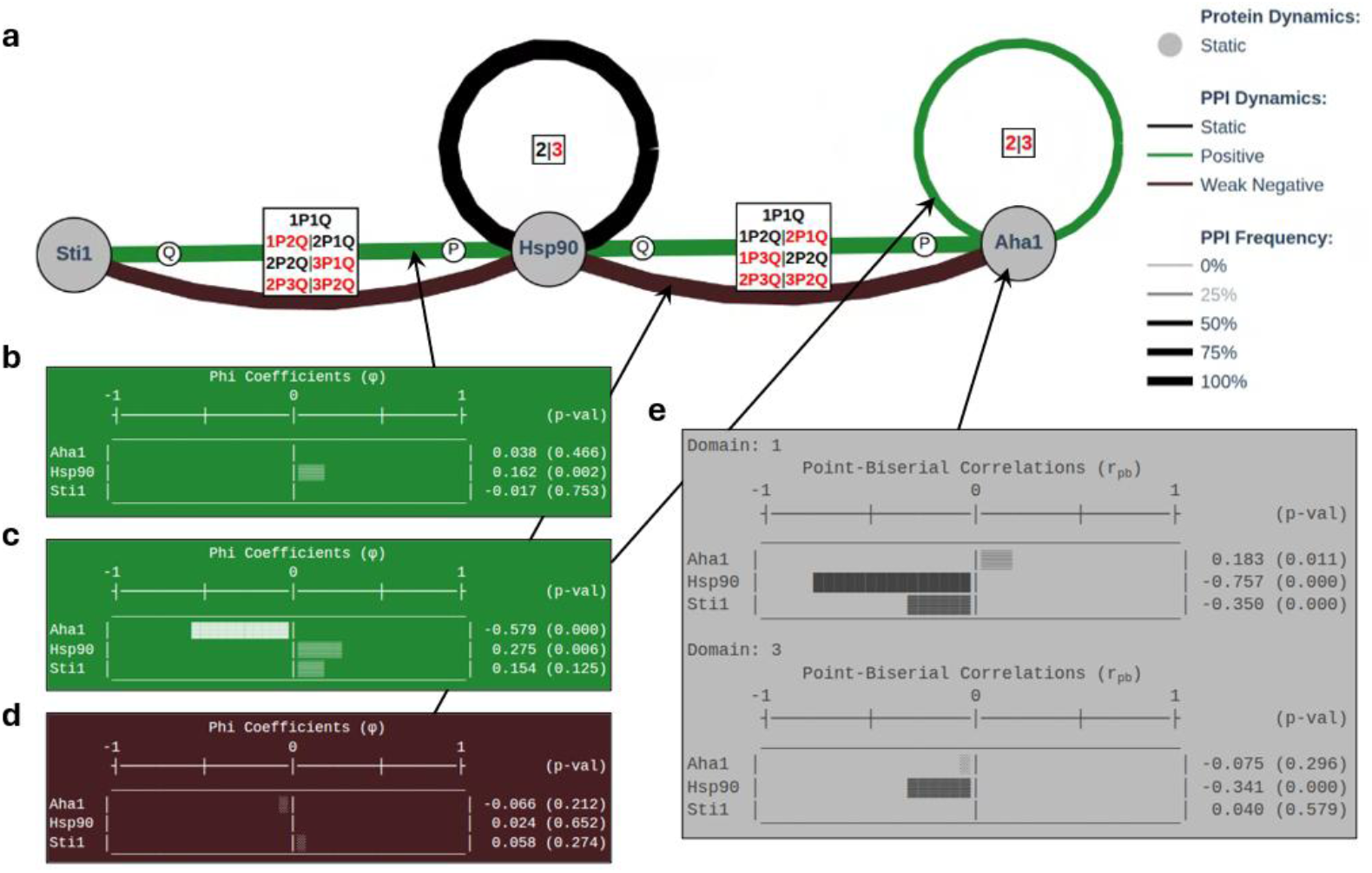
Interactive PPI graphs. PPI graph example of chaperone Hsp90 and the co-chaperones Sti1 and Aha1 system from *T. brucei*. (**a**) Resulting PPI graph. Insets show the identified stability of homooligomeric and multivalent heterooligomeric subassemblies. Circles with the letters P and Q indicate the correspondence of the stoichiometric coefficients from the insets. Bar plots showing the *φ* coefficients of the positive dynamic PPI between Sti1 and Hsp90 (**b**), the positive dynamic homooligomerization of Aha1 (**c**) and the weak dynamic negative PPI between Aha1 and Hsp90 (**d**). p-values are shown between brackets. (**e**) Bar plots with *r*_*pb*_ coefficients for the structured domains of Aha1.

#### 8.2. Association Between PPI Dynamics and Partners (φ coefficient)

To analyze the statistical significance of dynamic PPIs, MultimerMapper conducts a relationship analysis that associates the presence or absence of each possible protein partner with the presence or absence of each possible PPI within the predictions. For this, a phi-coefficient (φ) and its statistical significance using the *χ*^2^ test is computed [40]. The φ coefficient is built in a way that values significantly higher than zero (a p-value is provided) for a particular partner means that the partner is positively associated with the PPI. In other words, its presence is associated with the engagement of that particular PPI, and high φ values indicates that the correlation is strong. On the other hand, when this value is significantly smaller than zero, its presence is negatively associated with the engagement. This means that its presence is associated with the disruption of the PPI.

This data is included in the form of bar plots on each PPI mode metadata (**Fig. *8*b-d**). For the chaperone example, one of the PPI modes between Sti1 and Hsp90 is positively associated with Hsp90 presence (**Fig. *8*b**), as this PPI only appears when the models contain multiple subunits of Hsp90 (*i*.*e*., the dimer). Something similar happens for the homooligomerization of Aha1, being positively associated with Hsp90 presence **Fig. *8*c**, as it only appears when there are two subunits of Hsp90. Notice that Aha1 is negatively associated with its homointeraction. This is due to the fact that models that only contain Aha1 do not contain the PPI. Also, the φ value for Sti1 ispositive, but itsp-value is not significant. On the other hand, one of the PPI modes between Aha1 and Hsp90 is classified as weak dynamic negative, but all the associations are not significant (**Fig. *8*d**).

#### 8.3. Association Between Domain Conformation and Partners (r_pb_ coefficient)

Besides associating protein partners with predicted PPI dynamics, MultimerMapper also correlates the presence of specific partners with variations in RMSD using the point biserial coefficient (r_pb_). This correlation is performed only for well-structured domains (by default, a mean pLDDT > 60 in the reference structure) excluding disordered loops from the RMSD calculations (pLDDT < 70). Similarly to the φ coefficient, it varies between −1 and 1, with r_pb_ values significantly bigger than zero indicating a correlation between increase in RMSD (with respect to the reference domain structure) and the partner. Absolute values closer to 1 indicate a strong correlation and negative values are interpreted in the opposite way (lower of the RMSD).

For example, Aha1 has two structured domains, and MultimerMapper detects that both suffer conformational changes that lowers their RMSD that are associated with Hsp90 (**Fig. *8*e**). This association reflect conformational changes induced in Aha1 domains upon binding to Hsp90. Together, φ and r_pb_ coefficients aids in the identification of competitive binding, inhibition and activation of PPIs scenarios, through the association of changes in PPI patterns and conformational changes in the core segments of protein domains with specific protein partners. Potentially, these conformational changes can be induced by these partners, giving the users an intuitive and systematic way to obtain bioinformatic evidence with statistical support of functional mechanisms using conformational sampling coming from different modelling contexts.

## Discussion

This work presents a new way to systematically analyze predicted structures generated using deep learning structural models. It relies on the statistical analysis of associated confidence metrics and atomic coordinates to predict interactions and context-dependent conformational changes. The methodology is based on the systematic exploration of the stoichiometric space using an algorithm that seeks to balance the use of computational resources while maximizing the possibility of finding variations in protein interaction patterns and conformational states. In turn, the algorithm deciphers which protein combinations produce complexes where all the predicted subunits interact with each other, classifying them as “stable”. By adding new subunits to these stable combinations, we can discern those pathways that produce unstable combinations and discard them, reducing the exploration space. Ultimately, it is possible to identify the terminal combinations of these branches, which in theory have no further possible interaction surfaces with any of the protein entities in the system. This gives rise to the combinations we call “convergent stoichiometries,” which typically coincide with the stoichiometries reported in the literature. Furthermore, if the system contains multiple convergent stoichiometries, MultimerMapper is able to detect them.

However, there are systems where interaction surfaces give rise to symmetric complexes to which AlphaFold continues to add subunits beyond the expected point, resulting in convergent stoichiometries that differ from the reported stoichiometry. To overcome this, MultimerMapper implements other strategies that can be useful to discern between potential stoichiometries. These are the symmetry fallback detection and the pairwise confidence metric profiles. Either way, MultimerMapper’s predictions can only be as good as the deep learning model’s predictions. Regarding this, there is a direct limitation on the size of the convergent stoichiometries that can be detected, which is approximately 5000 amino acids within the AF3 server limits [18]. In turn, systems containing too many protein entities, while theoretically possible, cannot be directly analyzed because the number of predictions required is directly proportional to the square of the number of entities, resulting in high computational costs. Therefore, it is advisable to use as initialization system a set of protein entities with high interacting probability (*e*.*g*., from a Co-IP experiment). Nonetheless, the trends in deep learning structural models evolution indicates that they will become increasingly efficient, making it possible to model larger systems more quickly.

On the other hand, MultimerMapper is capable of constructing pseudo-trajectories based on the conformational states of individual proteins and their domains. While there is still debate as to whether conformational sampling of deep learning generated structures is truly representative of the conformational changes that proteins undergo [30, 41], these are representations of their conformational variability and can be a useful tool in certain situations. However, the methodology implemented by MultimerMapper from which conformational sampling is generated is fundamentally different from what has been described so far [42, 43]. Previous methodologies rely on sampling the input MSA, eliminating coevolutionary information that is ultimately reflected in the loss of predicted contacts, increasing the conformational variability of individual monomers. In contrast, MultimerMapper bases its exploration on contextual sampling, that is, systematically varying the subunits present in the predictions, without explicitly resorting to the elimination of coevolutionary information from the input MSA. This novel application allows for the discovery of correlations between the presence of certain proteins or combinations of them and the conformational states and confidence metrics of the different entities. These variations not only provide further evidence of PPI engagement but can be associated with protein functionality in the context of multiprotein complexes, being of great value for large dynamic systems where molecular dynamics is computationally infeasible when the objective is to analyze protein behavior in multiple contexts. In this sense, MultimerMapper manages to systematize this, solving it at a low computational cost.

Because MultimerMapper’s sampling algorithm was designed to produce suggestions that reduce computational cost, the number of chains of each protein entity present in the overall predictions is not the same. This has a clear implication for the interpretation of negative BPD values, since protein entities that are underexplored (for example, because they did not bind to any protein and therefore no N-mers containing them were generated) will tend to present negative values, making their interpretation difficult. In this sense, BPD values close to −1 could only be validly associated with the absence of a protein if, at some point in the pseudotrajectory, BPD values reach clearly elevated and sustained positive values. Therefore, it would be necessary to find an appropriate statistical test that would define the degree of significance of BPD values and identify those where BPD ≠ 0.

Although the methodology currently only considers polypeptide entities, it is feasible to apply it to other molecular entities. For example, a metric analogous to BPD could be generated, but one that measures the bias of small molecules or ions, associating their presence or absence with the conformational state of the different molecular entities in the systems. For example, by predicting the structure of guanine quadruplex (G4) forming DNA sequences using different numbers of potassium ions, we have successfully observed how these ions are associated with G4 conformations [44].

Another potential application is its use in the rational design of binders designed to disrupt interactions [45]. When designing binders targeting a protein surface that is intended to compete with an essential PPI, for example, of a pathogenic microorganism or a protein complex deregulated in tumorigenesis, MultimerMapper could be used to screen for designs with negative φ coefficients. That is, in addition to the usual screening measures for choosing potential synthetic designs (such as low PAE or low RMSD), this parameter could be included, keeping only those with predicted disruption capabilities.

Regarding PPI dynamics, it is worth mentioning that we observed a tendency in subsystems with expected homodimeric stoichiometries to not manifest the homointeraction PPI in the homotrimeric N-mer. That is, all subunits of the homotrimer are modeled as disconnected chains. As a consequence, PPIs are classified as negative dynamics, which does not seem to have a relevant biological interpretation. We think this is due to AF being unable to decide which pair of subunits to connect to assemble the dimer, unsuccessfully attempting to assemble a trimer instead. Another relevant related thing is that there may be situations where the PPI of a 2-mer that can form homo- or multivalent hetero-oligomeric structures larger than the 2-mer results in a false negative. Therefore, MultimerMapper will not continue exploring combinations involving that interaction. In these cases, it is possible to manually incorporate N-mers combinations of these proteins, since the PPI will sometimes manifest in more complex stoichiometries. If the PPI is detected and any N-mer is classified as stable, MultimerMapper will continue generating suggestions based on the stable stoichiometries. This results in a dynamic positive PPI that may or may not be relevant from a mechanistic point of view, as it can be just the product of a false negative.

With respect to multiple binding modes between protein pairs, MultimerMapper conducts the clustering algorithm only if it has predicted multivalency. This requires the simultaneous presence of more than one binding surfaces with the opposite partner [25], in one or both proteins, in a fraction of the input models. If a pair is considered monovalent, the clustering is not conducted, and all available contact matrixes are considered as a single cluster. While the problem has been solved for multivalent pairs, multiple binding modes are not always linked to multivalency. For example, some proteins can interact with the same surface via multiple conformational states [46]. The main difficulty for these cases is how to distinguish between a single cluster of contact matrixes versus multiple ones. We are working to solve this situations by performing the hierarchical clustering using a cluster size of 2 and applying an RMSD threshold between the resulting representative structures. If these two potential binding modes differ significantly, the default matrix clustering algorithm can be triggered to identify the number of different binding modes. This solution will be incorporated into future versions of the software. From a practical perspective, the program was designed to assist in systematically interpreting structural predictions, targeting users without extensive experience in structural and computational biology. Furthermore, independence from third-party software was sought for ease of use. Therefore, each run of MultimerMapper generates an interactive HTML-based graphical report that can be run in any browser. It has integrated structure viewers and interactive graphs that allow for easy interpretation of the behavior of the system of interest (**Supplementary Figure 4**). For advanced users, the software can also be integrated into other bioinformatics pipelines using the various packages available in MultimerMapper.

In summary, MultimerMapper is a tool that implements a novel methodology that can be well adapted by the scientific community to understand the behavior of molecular systems involving the simultaneous interaction of multiple proteins. It does not require computer expertise or sophisticated structural tools, being particularly useful for analyzing completely unknown and highly dynamic systems, as it requires no prior information beyond the amino acid sequence. In future iterations, it will be possible to integrate other types of molecules with biological interest and evaluate their association with the conformational behavior of different biopolymers.

## Methods

### Protein Structure Prediction

Structure predictions were performed with AlphaFold3 server [18] and a local instance of AlphaFold2-multimer v3 [2]. For AF2m, we used as input the multiple sequence alignments (MSAs) described by Bryant et al. [3], without using templates. For this, the Discoba database [47] was adapted to allow sequence matching and improve alignment coverage within the trypanosomatid taxonomic group by concatenating the alignment generated from the modified database with the MSA generated by the ColabFold MSA server [4], as described by Wheeler et al. [47]. For each protein combination, 5 models were generated, using a maximum of 20 recycles, stopping recycling if two successive models reached a ΔRMSD < 0.5 Å. The runs were executed within Amazon Web Services (AWS) using g4dn.12xlarge, g5.48xlarge, p3.2xlarge, p3.16xlarge, or p3dn.24xlarge instances, depending on the dataset size and the total residues of individual predictions. For a full description of the Discoba database tuning and MSA generation, see the supplementary material.

### Models Pairwise Decomposition and Metrics Extraction

Each model of *N* protein chains (*h*_1_, *h*_2_, …, *h*_*N*_) was decomposed into all possible paired combinations. To do this, for each chain pair *h*_*p*_, *h*_*q*_, the combined atomic coordinates of *h*_*p*_ and *h*_*q*_ were extracted into a sub-model, along with their per residue pLDDT values, and the two inter-chain off-diagonal submatrices of the model’s PAE matrix corresponding to residue pairs *i, j* of the chain *h*_*p*_ → *h*_*q*_ and *h*_*q*_ → *h*_*p*_, respectively. One of the submatrices was transposed, and the minimum inter-chain PAE matrix was calculated by assigning the minimum observed value for each pair *i, j*. The overall minimum inter-chain PAE value (miPAE) was then calculated by taking the smallest element of the resulting matrix. The predicted docking coefficient (pDockQ) was algo computed [3]. In addition, the model’s pTM and ipTM values were extracted, ranking them based on the ipTM value relative to the other models within each prediction, assigning rank = 1 to the model with the highest ipTM. In the case of 2-mers, the models were not split into sub-models, as they represented pairwise interactions directly. For 2-mers, the same metrics as for N-mers were calculated.

For each sub-model (or 2-mer), the identifier corresponding to each protein, the identifiers of each chain, the protein composition of the original model, the sub-model atomic coordinates, the minimum inter-chain sub-PAE matrix, rank, pTM, ipTM, miPAE, and pDockQ were stored in tabular format for further processing.

### Protein-Protein Interaction and Residue-Residue Contact Prediction

PPI detection was performed for each pair of protein entities *P, Q* within each prediction composed of 5 models that had at least one chain of *P* and one chain of *Q*. By default for 2-mers, the PPI between *P* and *Q* was considered positive in a prediction if at least 4 models had *miPAE* ≤ *miPAE*_*cutoff*_ (AF2m: *miPAE*_*cutoff*_ = 10 Å for *FPR* ≅ 0.05 and *TPR* ≅ 0.58 at *N*_*models*_ = 4 /AF3: *miPAE*_*cutoff*_ = 8.3 Å for *FPR* ≅ 0.05 and *TPR* ≅ 0.58 at *N*_*models*_ = 4). For N-mers, the PPI between *P* and *Q* was considered positive if at least 4 models had at least one pair of chains of *P* and *Q* (decomposed sub-models) that exceeded the established miPAE and *N*_*models*_ cutoff values.

In cases where no PPI was detected for a pair within a prediction, all *P, Q* sub-models within the same prediction were considered to have no residue-residue contacts (RRCs). Otherwise, the center of mass per residue (centroid) of each chain in each submodel was calculated and an intercentroid distance matrix *D* of dimensions *L*_*P*_ × *L*_*Q*_ was constructed, where *L*_*P*_ and *L*_*Q*_ are the sequence lengths of *P* and *Q*, respectively. RRCs were considered to be those pairs of residues *i, j* with intercentroid distances *D*_*ij*_ ≤ 8 Å and *miPAE*_*ij*_ ≤ *miPAE*_*cutoff*_. The RRCs of each sub-model were represented in a contact matrix of dimensions *L*_*P*_ × *L*_*Q*_, where 1 or 0 was assigned to each position *i, j*, depending on whether the residue pair was in contact or not, respectively.

### Protein Segmentation and Domain Detection

The segmentation of a protein *P* of sequence length *L*_*P*_ into compact and well-structured segments separated by disordered segments was carried out by processing the intra-chain PAE matrix of its reference structure with a graph-based community clustering algorithm, modified from the algorithm described in https://github.com/tristanic/pae_to_domains. First, the reference structure was defined as the chain of *P* with the highest average pLDDT. Then, a graph with *L*_*P*_ vertices (one per residue) was constructed and only the residue pairs *i, j* with *PAE*_*ij*_ < 5 Å were connected with edges, assigning each pair a weight *w*_*ij*_ = 1/*PAE*_*ij*_. Residues belonging to the same communities were identified using the Laiden clustering algorithm [48], controlling the granularity with a resolution parameter *g*_*r*_ scaled in 1/100 (smaller *g*_*r*_ → fewer and larger domains /larger *g*_*r*_ → more and smaller domains). Finally, residue clusters from a small community (usually corresponding to disordered loops) surrounded by a larger community were integrated into the larger community.

To fine-tune domain assignment using the *g*_*r*_ value, 2D visualizations were generated with the intrachain PAE matrix heat map and 3D visualizations representing the protein carbon backbone, both showing the assigned domains with different colors. This allowed interactively assigning the best *g*_*r*_ values for each protein entity (**Supplementary Figure 3**).

### RMSD Pseudo-Trajectories and Bias in Partners Distribution (BPD)

For each protein entity *P* present in a set of *c* predictions with different combinations of *n* protein entities, RMSD pseudo-trajectories were constructed as described below. All atomic coordinates of all chains of *P* present in the *c* predictions were isolated, resulting in a list of *m* isolated models (*P*_1_, *P*_2_, …, *P*_*m*_), registering the stoichiometric coefficient of each protein entity present in the original combination. The reference structure (*P*_*ref*_) was identified as the model with the highest average pLDDT, the atomic coordinates of the *α*-carbons of each model *P*_*i*_ were structurally aligned with those of *P*_*ref*_ using the Kabsch algorithm [49], and the *RMSD*_*i*_ value was calculated. The *m* models were sorted by ascending RMSD values, resulting in a pseudo-trajectory of *m* frames sorted by structural similarity. The root mean square fluctuation (RMSF) per residue of the pseudo-trajectory was calculated. For each frame *j*, the pLDDT value per residue, the average pLDDT value, and the radius of gyration (ROG) were recorded/computed. In addition, the Bias in Partner Distribution (BPD) was calculated for each position *j* and each of the *n* protein entities. This parameter measures how locally enriched (*BPD* ≅ 1) or depleted (*BPD* ≅ −1) a partner *k* is along the pseudo-trajectory with respect to its global frequency *f*^*global*^. To do this, an *n* × *m* boolean matrix was constructed, recording for each *j*-th frame whether the *k*-th protein entity was present or not in the original prediction. Thus, for each protein entity *k*, its *f*^*global*^ was calculated as

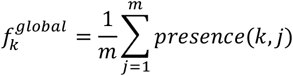

where

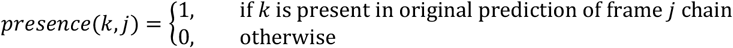

In the case where *k* = *P, k* was considered present only if *k* was found in at least two units. On the other hand, the local frequency (*f*^*local*^) was calculated in a rolling and centered manner for different window values *w* ∈ {5,10,15,20} as

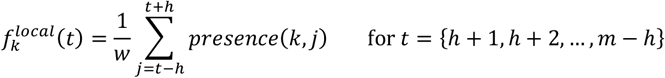

where *h* = ⌊*w*/2⌋ and *t* is the frame adjusted to the window size so as not to exceed the range *j* ∈ [1, *m*]. In this way, the BPD was defined for each frame t as

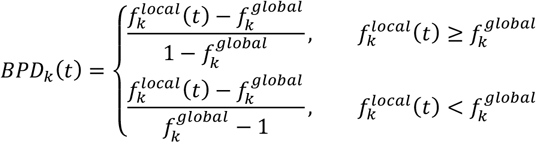

With all the collected information, each BPD_k_, the RMSD, the pLDDT per residue in the form of a heat map, the average pLDDT and the ROG as a function of frame *j* were plotted, adding gaussian noise to the BPD to avoid overlapping of the curves. A final structural alignment step was applied to smooth out alignment fluctuations due to disordered loops. Only the *α*-carbons of residues with the highest median pLDDT (consistently well predicted) and lowest interquartile range (low variability) were used simultaneously (see supplementary material for a detailed description of how to identify core residues based on pLDDT to perform the final alignment).

If more than one domain segment was assigned to *P* by the domain detection algorithm, the same algorithm was run for each segment but restricted to the residues and atoms within that segment, assigning a new reference structure for each domain.

### pLDDT Clustering and Partners Enrichment Analysis

For each protein entity *P* of *L*_*P*_ residues in length present in a group of *c* predictions with different combinations of *n* protein entities, the per residue pLDDT profiles of *P* were clustered according to their similarity, and the partner enrichment was analyzed within each group, as described below.

All atomic coordinates of all *P* chains present in the *c* predictions were isolated, resulting in a list of *m* models (*P*_1_, *P*_2_, …, *P*_*m*_) and the stoichiometric coefficient of each protein entity present in the combination of origin was recorded. Let *X* = [*pLDDT*_*r,j*_] be an *m* × *L*_*P*_ matrix, where *m* is the number of models and *pLDDT*_*r,j*_ the pLDDT value of residue *r* of the *j*-th model. Clustering between 2 and *K*_*max*_ = min (10, *m*) was explored using the K-means algorithm on the matrix *X*, distributing the *m* models into K groups. For each K evaluated, its average silhouette score 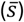 was calculated, and the *k* value that maximized 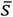 was chosen. For each group, the average pLDDT per residue was plotted as a solid line and its standard deviation as a transparent band.

For partner enrichment, the stoichiometric coefficients of the protein entities associated with each model were analyzed, and a table of partner frequencies was constructed within each group k. For each protein partner *Q*, its frequency *f* in group *k* is given by

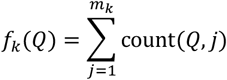

where *m*_*k*_ is the number of models in group *k* and *count*(*Q, j*) is the number of times *Q* appears in model *j*. If *Q* = *P*, one unit was subtracted from *count*(*Q, j*) to avoid counting itself. Finally, the frequencies within each *k* were represented as a word cloud, where the size of each *Q* name is directly proportional to *f*_*k*_(*Q*).

### Multivalency Detection

Let *P* and *Q* be two identical or distinct protein entities that establish PPI in at least one of all 2-mer or N-mer combination predictions, their multivalent interactions were determined using the multivalent chains fraction (MCF). To do this, all decomposed models of *P* and *Q* coming from N-mers that contained 3 subunits of *P* and *Q* (1P2Q or 2P1Q) were taken, regardless of the presence or absence of other proteins. Then, for each matching N-mer, the number of *Q* chains in contact with each *P* chain was counted, and vice versa. Two chains were considered in contact if they had at least 5 RRCs (see RRC prediction method). Multivalent chains (*C*_2_) were considered to be those in contact with ≥2 chains of the opposite protein entity, and monovalent chains (*C*_1_) were considered to be those in contact with ≤1 chain of the opposite protein entity. The MCF was defined as

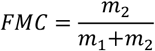

where *m*_1_ and *m*_2_ are the number of observations for *C*_1_ and *C*_2_, respectively. The *P*-*Q* pair was considered multivalent if *FMC* ≥ 0.2. In cases where *FMC* < 0.2 and N-mers with more than 3 combined subunits of *P* and *Q* were available, it was recalculated using all N-mers and the *P*-*Q* pair was reconsidered using the same *FMC* cutoff.

### Contact Matrix Clustering Algorithm

Let *P* and *Q* be two protein entities of sequence length *L*_*P*_ and *L*_*Q*_, respectively, which can be identical or different, and interact with each other in a multivalent manner. Let *M* be a list of *m* Boolean contact matrices between *P* and *Q* (*M* = {*M*_1_, *M*_2_, …, *M*_*m*_}) from the RRC detection algorithm (see above) and *T* the list of decomposed sub-structures of *P* and *Q* that gave rise to the matrices *M* (*T* = {*T*_1_, *T*_2_, …, *T*_*m*_}). The clustering algorithm described below was applied.

A distance matrix *D* between the *m* models was calculated from the RMSD values of the residues involved in contact in at least one of the matrices of each pair. To do this, for each pair of decomposed models *u, v*, the lists of residues involved in contacts (*Ω*) of *P* and *Q* were defined as

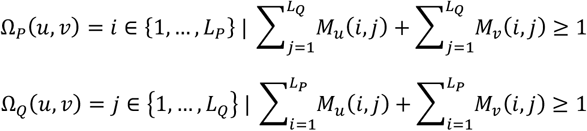

That is, *Ω*_*P*_ and *Ω*_*Q*_ are the indices of residues *i* and *j* of *P* and *Q*, respectively, that are in contact in at least one of the two matrices. Atomic coordinates of the *α*-carbons of *Ω*_*P*_ and *Ω*_*Q*_ residues of each *T*_*u*_, *T*_*v*_ pair were extracted, and the atomic coordinate lists *xyz*_*u*_(*u, v*) and *xyz*_*v*_(*u, v*) were constructed. Both lists have the same length for each *u, v* pair and are sorted in ascending order by index in *Ω*, concatenating first those residues of *Ω*_*P*_ and then those of *Ω*_*Q*_, maintaining an element-by-element correspondence. The *xyz*_*v*_(*u, v*) coordinates were aligned to those of *xyz*_*u*_(*u, v*) using the Kabsch algorithm [49], and the RMSD was calculated. The process was repeated for each *u, v* pair, and the *m* × *m*-dimensional distance matrix *D* was constructed from the results.

*D* was used as input distance matrix for the bottom-up hierarchical clustering algorithm using the average linkage method. The number of clusters to explore was set between *K*_*min*_ and *K*_*max*_, which depend on the maximum valency (*ψ*) of the chains observed during the multivalency detection algorithm (see above) and the number of matrices to be clustered (*m*), as follows:

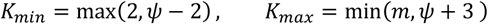

For each cluster number *K*, the average silhouette coefficient 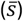 was calculated from the resulting groups and the distance matrix, choosing the optimal number of clusters (*K*_*op*_) as the one that maximizes 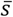.

Finally, the contact matrices for each cluster *K* were averaged to create the average contact matrices 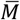, which represent relative frequency of contacts by position *i, j*. That is, the one calculated as

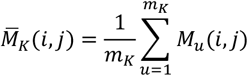

where *m*_*K*_ is the number of matrices in cluster *K*. A representative structure was taken for each cluster by taking the model whose contact matrix had the smallest Jaccard distance (intersection over union) to the binarized average matrix.

To construct the MDS visualizations, *D* was passed as argument of the MDS function of sklearn package, setting two as number of components. The resulting coordinate components of each matrix were plotted in a dispersion plot, coloring them by cluster.

### Stability, Convergence and Stoichiometry Inference

#### N-mer Stability Definition

An N-mer composed of *N* protein chains (*h*_1_, *h*_2_, …, *h*_*N*_) with five available structural models was defined as stable if enough models produced fully connected inter-chain contact graphs independently. The definition is described below.

Each model of rank *r* (ipTM rank) was converted into an undirected graph (*G*_*r*_), representing the chains as nodes and connecting with edges those *h*_*p*_, *h*_*q*_ pairs that had a number of contacts *C* between all their interchain residue pairs *i, j* that exceeded a cutoff value (*C*_*cutoff*_ = 5 by default). That is, nodes *h*_*p*_ and *h*_*q*_ were connected when

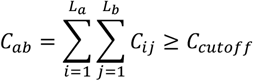

where *L*_*p*_ and *L*_*q*_ are the sequence lengths of *h*_*p*_ and *h*_*q*_, respectively; and *C*_*ij*_ is given by

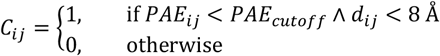

where *PAE*_*ij*_ and *d*_*ij*_ are the inter-chain PAE and inter-centroid distance values, respectively, for the residue pair *i, j*; and *PAE*_*cutoff*_ is the PAE cutoff value (*PAE*_*cutoff*_ = 8.3 Å by default). *G*_*r*_ was considered fully connected if, for every possible pair of nodes, there is a path between them. Next, the number of *G*_*r*_ fully connected (*R*) was quantified, considering the N-mer stable if *R* ≥ *r*_*cutoff*_ (*r*_*cutoff*_ = 3 by default). Otherwise, it was defined as unstable.

If *N* = 2 (2-mers), the combination was considered stable if it exceeded the PPI detection cutoff (see PPI detection method).

#### Stoichiometric Space Exploration Algorithm and System Convergence

Each system of *n* protein entities was analyzed using a pipeline consisting of rounds of protein structure prediction, analysis (PPI prediction, RRC prediction, and detection of stable N-mers), and generation of combinations (suggested N-mers) of the *n* entities based on observations during the analysis section. At the start of each new round, the suggested predictions were added to the predictions from the previous round, entering them into a new analysis stage.

Systems were initialized by predicting all possible pairwise combinations of the *n* entities (2-mers), including homodimers. After detecting interacting 2-mers (see PPI prediction method), *n* - *f* suggestions of size *N* = 3 (3-mers) were generated for each stable 2-mer by adding one unit of each possible protein entity involved in PPIs to the base dimeric combination, where *f* is the number of protein entities not involved in PPIs. The system was represented as a directed graph, where the resulting stoichiometries (suggested combinations) were considered children of the stable 2-mers that gave them origin. In this graph, each daughter could have more than one parent, and unstable 2-mers did not generate daughter stoichiometries.

The suggestions were predicted and incorporated into a new round of analysis with the 2-mers. In this new round, stable 3-mers were detected and *n* − *f* new suggestions of *N* = 4 were generated for each stable 3-mer, without generating children for unstable 3-mers. The process was repeated, increasing *N* + 1 in each iteration, expanding the graph of parent and daughter stoichiometries per iteration. The process stopped when the system reached convergence, that is, when no suggested stoichiometries remained unpredicted. Convergent stoichiometries of the system were identified as those stable stoichiometries where all of their children were unstable and none of their possible children were left unexplored.

For systems where all *N* = 3 suggestions were unstable (e.g., nucleosome), in addition to the *N* + 1 increments, *N* + 2 increments were incorporated for each iteration. To do this, *d* extra suggestions were generated for each stable 2/N-mer by adding the stoichiometry of each possible stable 2-mer to the stoichiometry of the base stable 2/N-mer, where *d* is the number of stable 2-mers. *N* + 2 suggestions were considered children of the N-level stoichiometries that gave rise to them only if no stable intermediate (*N* + 1) stoichiometry that could have given rise to them were found.

### Symmetry Fallback Detection

Let *P* be a homooligomerizing protein entity and *Z* − 1 successive homooligomeric predictions of *P* are available, ranging from the dimer (homo-2-mer) to the homooligomer of size Z (homo-Z-mer). The presence of a symmetry fallback was defined by detecting a drop greater than 20% in the estimated radius of the top-ranked model (ipTM rank) in any homo-N-mer (*R*_*N*_) relative to the estimated radius of its predecessor top-ranked model (*R*_*N*−1_). To estimate the radius, for each chain *h* of each rank 1 model, the center of mass (CM) of the domains involved in homooligomerization (*D*) was computed according to

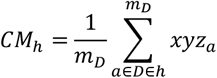

where the sum is performed over the atoms (*a*) of *D* of chain *h, m*_*D*_ being the number of atoms in *D*, and *xyz* being their atomic coordinates. The global center of mass of the model was then calculated, restricting it to the domains involved in homooligomerization as

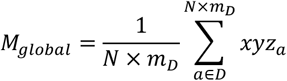

The Euclidean distance (*d*) of each chain to the *CM*_*global*_ was calculated, estimating *R*_*N*_ with the average of the Euclidean distances 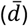:

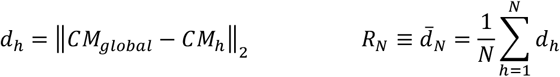

In cases where a symmetry fallback was detected for an N-mer (*R*_*N*_ < 0.8 × *R*_*N*−1_), it was determined which stoichiometry it resembled by comparing its radius (*R*_*F*_) with that of N-mers predecessors (*R*_*N*<*F*_). For this purpose, a 95% Student’s T confidence interval with *N* − 1 degrees of freedom (CI) of *R*_*F*_ was constructed and it was checked whether any *R*_*N*<*F*_ was contained by CI. The first homooligomer (in decreasing N size order) of size *T* whose *R*_*T*_ was contained by CI was assigned as the falling back stoichiometry, assigning the nomenclature *F*_*f*_*T* to indicate that the homooligomer of size *F* undergoes a symmetry fallback that makes it similar to the homooligomer of size *T* (*F* falls back to *T*). If no *R*_*N*<*F*_ was contained by CI, the radii were reestimated using a two-dimensional projection (see supplementary material for a description of the PCA projection) and the process was repeated using the CI of the projected radius and the projected radii. In case neither of the two methods covers the R of a smaller homo-N-mer, the *R*_*T*_ that minimizes the difference |*R*_*F*_ − *R*_*N*<*F*_| was chosen.

### Correlation Between PPI Dynamics and Protein Presence Using Phi Coefficient

The tendency to stabilize or destabilize a PPI between a pair of protein entities *P* and *Q* induced by the presence of a third protein entity *S* was quantified using the phi correlation coefficient (*φ*). To do this, all decomposed models of *P* and *Q* derived from N-mers were analyzed, and a 2 × 2 matrix was constructed representing the count of detection (1) or non-detection (0) of PPIs in the columns, and the count of presence (1) or absence (0) of *S* in the model in the rows. The coefficient *φ* was computed as

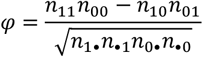

where *n*_11_ is the number of decomposed models in which a PPI was detected and *S* was present; *n*_00_ is the number of those in whom PPI was not detected and *S* was not present; *n*_10_ is the number of those in whom PPI was detected and *S* was not present; *n*_01_ is the number in whom PPI was not detected and *S* was present; and *n*_1•_, *n*_•1_, *n*_0•_, and *n*_•0_ are the marginal totals.

In cases where *P* ≠ *Q* and *S* ∈ {*P, Q*}, *S* was considered present in the model only if it was found in ≥2 units. For cases where *P* = *Q* = *S, S* was considered present in the model only if it was found in ≥3 units. PPI detection was defined as the presence of at least 5 residue-residue contacts between *P* and *Q*. The null hypothesis was no association between the presence of *S* and PPI detection (*H*_0_: *φ* = 0), and the alternative hypothesis was association (*H*_1_: *φ* ≠ 0), which was evaluated using the Chi-Square (*χ*^2^) test. Values of *φ* > 0 (p-val < 0.05) were associated with a positive correlation between the presence of *S* and the establishment of PPI, while values of *φ* < 0 (p-val < 0.05) were considered a negative association.

### Correlation Between RMSD and Protein Presence Using Point-Biserial Correlation

The correlation between the conformational state of an ordered domain *D* in a protein entity *P* and the presence of another protein entity *S* was quantified using the point-biserial correlation coefficient (*r*_*pb*_). To do this, the segment corresponding to *D* was isolated from all *m* isolated chains of *P*, and the root mean square deviation (RMSD) value was calculated against the segment *D* of the reference structure. An *m* × 2 table was then constructed with the RMSD values in one column and the presence (1) or absence (0) of *S* in the model from which the isolated chain originated in the other. The coefficient *r*_*pb*_ was computed as

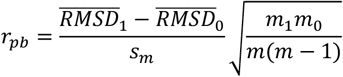

where *s*_*m*_ is the standard deviation of all chains; 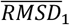 and 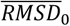 are the average RMSDs of the chains where *S* was present or absent, respectively; and *m*_1_ and *m*_0_ are the number of chains where *S* was present or absent, respectively.

Ordered domains of *P* were defined as those segments defined by the domain detection algorithm (see protein segmentation method) with *mean*(*pLDDT*) > 60 values in the reference structure. Only the *α*-carbons of residues with *pLDDT* > 70 were used to calculate the RMSD. In cases where *P* = *S, S* was considered present in the model only if found in ≥2 units. The null hypothesis was the absence of an association between the RMSD value and the presence of *S* (*H*_0_: *r*_*pb*_ = 0), and the alternative hypothesis was the association (*H*_1_: *r*_*pb*_ ≠ 0), which was tested using a T-test with *m* − 2 degrees of freedom. Values of *r*_*pb*_ > 0 (p-val < 0.05) were interpreted as a positive association (the presence of *S* is associated with RMSD values further from the reference) and values of *r*_*pb*_ < 0 (p-val < 0.05) as a negative association (RMSD values closer to the reference).

### Pairwise Confidence Profiles

Let *P* and *Q* be two protein entities, which may be identical or distinct, that were detected as interactors using the PPI detection method (see PPI prediction method), each present in at least one subunit in *c* structural predictions. Each structural prediction consists of five models and was generated using different combinations of *n* protein entities. The construction of confidence profiles for the *P*-*Q* pair is described below.

The pLDDT values per residue were extracted from all *P* and *Q* chains, and the resulting values were grouped according to the structural prediction (combination of proteins) and model number. For each group, the average pLDDT, its standard deviation (SD), median, first quartile (Q1), and third quartile (Q3) were calculated.

From the PAE matrix of each model, the two PAE submatrices corresponding to the interchain residues of each pair of *P* and *Q* chains were extracted. The two matrices were combined by transposing one of them and maintaining the minimum value for each residue pair. The average PAE value of each resulting matrix (aiPAE) was then calculated, and the miPAE values were extracted. The aiPAE and miPAE values were grouped by structural prediction and model, and the mean, SD, median, Q1, and Q3 were calculated for each group.

The pLDDT, aiPAE, and miPAE profiles were constructed by plotting the average pLDDT and its SD (or median and interquartile range) as a function of model number and colored according to the protein combination of each structural prediction.

## Data Availability

The source code of MultimerMapper can be found at https://github.com/elviorodriguez/MultimerMapper. The code used to generate the sequence alignments and protein structure predictions with AlphaFold2-multimer and the modified Discoba database can be found at https://github.com/elviorodriguez/DiscobaMultimer.

## Acknowledgements

This work was funded by AWS Cloud Credit for Research Program (PS_R_FY2022_Q2_CONICET_Serra_10900), the National Research Council (CONICET: PIP 2021-0848), the National Agency for Science (ANPCyT: 2020-01704), and Universidad Nacional de Rosario (UNR: 0001-00285485). We want to thanks Juan Manuel Faus Gonzáles, Valentina Nora Faus Gonzáles, Damian Molini and Gustavo Guerrero Chin-Aleong for developing and implementing an interface to organize and access the structural predictions generated for this article. We also thanks VEuPathDB for providing access to essential bioinformatics resources and databases that facilitated our research.

